# Global and regional ecological boundaries drive abrupt changes in avian frugivory interactions

**DOI:** 10.1101/2021.09.18.460873

**Authors:** Lucas P. Martins, Daniel B. Stouffer, Pedro G. Blendinger, Katrin Böhning-Gaese, Galo Buitrón-Jurado, Marta Correia, José Miguel Costa, D. Matthias Dehling, Camila I. Donatti, Carine Emer, Mauro Galetti, Ruben Heleno, Pedro Jordano, Ícaro Menezes, José Carlos Morante-Filho, Marcia C. Muñoz, Eike Lena Neuschulz, Marco Aurélio Pizo, Marta Quitián, Roman A. Ruggera, Francisco Saavedra, Vinicio Santillán, Matthias Schleuning, Luís Pascoal da Silva, Fernanda Ribeiro da Silva, Sérgio Timóteo, Anna Traveset, Maximilian G. R. Vollstädt, Jason M. Tylianakis

## Abstract

Species interactions can propagate disturbances across space, though ecological and biogeographic boundaries may limit this spread. We tested whether large-scale ecological boundaries (ecoregions and biomes) and human disturbance gradients increase dissimilarity among ecological networks, while accounting for background spatial and elevational effects and differences in network sampling. We assessed network dissimilarity patterns over a broad spatial scale, using 196 quantitative avian frugivory networks (encompassing 1,496 plant and 1,003 bird species) distributed across 67 ecoregions and 11 biomes. Dissimilarity in species and interactions, but not in network structure, increased significantly across ecoregion and biome boundaries and along human disturbance gradients. Our findings suggest that ecological boundaries contribute to maintaining the world’s biodiversity of interactions and mitigating the propagation of disturbances at large spatial scales.

**One-Sentence Summary:** Ecoregions and biomes delineate the large-scale distribution of plant-frugivore interactions.

## Main text

Abiotic gradients underlie the existence of a wide array of natural ecosystems, which are the cornerstone of biological diversity on Earth (*1, 2*). Ecoregion borders delineate regional discontinuities in the environment and in species composition (*3, 4*), whereas biomes mark these ‘break points’ at a global scale, such that ecoregions are nested within biomes (*1*, *4* and fig. S1). Accordingly, ecoregion and biome maps have been widely used for delimiting terrestrial ecosystems and guiding conservation planning (*4, 5*), but the question of whether distinct ecoregions truly represent sharp boundaries for species composition across several taxa was only recently answered on a global scale (*3*).

There has been growing recognition that interactions among species are critical for biodiversity and ecosystem functioning (*6*) and an important component of biodiversity in themselves, such that interactions may disappear well before the species involved (*7*). Species interactions also provide a pathway for the propagation of disturbances via direct and indirect effects, such as secondary extinctions and apparent competition (*8, 9*), potentially connecting species at a global scale. Thus, both natural and human disturbances in local communities of interacting species might reverberate and affect ecosystem functioning at multiple sites (*10, 11*). However, the spread of disturbances may be hindered when interactions are arranged into distinct compartments (*12*). Despite this importance, we are only beginning to understand whether such discontinuities exist in ecological networks at very large scales (*10, 11*), such as across ecoregions and biomes, potentially acting as a barrier to the global spread of disturbances. Because species tend to be replaced across ecosystems (*2, 3*) and environmental conditions can favour some types of interactions over others (*13*), we hypothesize that the large-scale distribution of species interactions is punctuated by ecoregion and biome boundaries. Alternatively, even though at smaller scales habitats may differ in their interactions (*14*), interactions that occur across habitat boundaries can connect their assemblages, causing multiple habitats to function as a single unit (*9*). Moreover, ecological boundaries might be further blurred by the processes of species and interaction homogenization, which accompany land-use change and biotic invasion (*10, 15*). Thus, an alternative hypothesis would be that shared interactions and biotic homogenization prevent any sharp discontinuities in interaction composition (i.e., the identity of interactions). Importantly, natural and human-disturbance gradients are juxtaposed with spatial processes that drive gradual changes in species and interaction composition (*13*). Indeed, distance-decay relationships have been demonstrated across spatial and elevational gradients not only for species (*16*), but also for ecological networks (*17–19*), and likely result from dispersal limitation and increasing environmental dissimilarity with increasing geographic distance (*16*).

Here we evaluate whether significant changes in the composition of species, the composition of interactions, and the structure of local networks of avian frugivory are driven by large-scale ecological boundaries (ecoregions and biomes) and human disturbance gradients, while accounting for background spatial and elevational effects. Given known patterns of species turnover across environmental gradients (*16*), we hypothesize a similar pattern of interaction dissimilarity, which could potentially lead to changes in the whole structure of networks. We focused on frugivory networks because of their importance for seed dispersal (*20*), promoting species diversity (*21*) and regenerating degraded ecosystems (*22*). As such, mapping the large-scale distribution of plant-frugivore interactions will be crucial to ensure ecosystem functioning and resilience in a context of increasing global changes.

To test our hypotheses, we assembled a large-scale database comprising 196 quantitative local networks of avian frugivory (with 9,819 links between 1,496 plant and 1,003 bird species) distributed across 67 ecoregions, 11 biomes, and 6 continents (figs. S1, S2 and table S1). To ensure that our results would not be driven by taxonomic uncertainty and sampling effects, we standardized the taxonomy of plant and bird species in our networks following a series of steps (figs. S3 to S6) and controlled statistically for network sampling metrics in our analyses [see materials and methods (*23*)].

**Table 1.** Multiple drivers of plant-frugivore interaction dissimilarity. Here, we used the binary version of ecoregion and biome distance matrices. *P* values were calculated using a combination of generalized additive models and multiple regression on distance matrices. EDF represents the effective degrees of freedom for each smooth term in the model. Bold values indicate statistically significant results (*P* < 0.05). *N* pairs of networks = 19,110.

| **Parametric coefficients** | **Estimate** | **t** | **p** |
| --- | --- | --- | --- |
| Intercept | 0.997 | 2964.191 | **0.001** |
| Ecoregion (same) | -0.070 | -36.401 | **0.001** |
| Biome (same) | -0.002 | -3.323 | **0.044** |
| **Smooth Terms** | **EDF** | **F** | **p** |
| s (local footprint distance) | 8.534 | 29.988 | **0.001** |
| s (spatial distance) | 8.785 | 65.378 | **0.001** |
| s (elevational difference) | 6.168 | 47.707 | **0.001** |
| s (hours distance) | 1.558 | 5.449 | 0.290 |
| s (months distance) | 5.482 | 6.902 | 0.075 |
| s (years distance) | 7.208 | 11.848 | **0.019** |
| s (sampling intensity distance) | 1.018 | 5.182 | 0.259 |
| s (methods distance) | 8.632 | 16.002 | **0.005** |

We generated several distance matrices (N × N, where N is the number of local networks in our dataset) to be our variables in the statistical models (*23*). Specifically, we used ecoregion, biome, local human disturbance [measured using the human footprint index (*24*)], spatial, elevation and sampling-related (i.e., hours, months, years, intensity and methods; figs. S7, S8 and table S2) distance matrices as predictor variables, and facets of network dissimilarity as the response variable. We constructed two distinct versions of the ecoregion and biome distance matrices: in the binary version, pairs of networks were given a dissimilarity of zero if they came from localities within the same ecoregion/biome, otherwise one; in the quantitative version, we calculated a continuous measure of environmental distance between the ecoregions and biomes where the networks were located (*23*). We used three response variables: species turnover (β_S_), which estimates the pairwise dissimilarity in species composition between networks (*25*), interaction dissimilarity (β_WN_), which represents the pairwise dissimilarity in the identity of interactions between networks (*25*), and network structural dissimilarity, which captures differences in the number of links in the networks, their relative weightings, and their arrangement among species. To generate this latter metric, we combined several network descriptors (weighted connectance, weighted nestedness, interaction evenness, PDI and modularity) using Principal Component Analysis (*23*). To evaluate the effect of each of our predictor distance matrices on our response variables, we employed a combination of Generalized Additive Models and Multiple Regression on Distance Matrices (*26*). Finally, we explored the unique and shared contributions of our predictor variables to network dissimilarity using deviance partitioning (*23*).

As expected based on the definition of ecoregions and biomes, the turnover of plant and frugivorous-bird species composition was strongly affected by ecoregion and biome boundaries (tables S4 and S5; fig. S9, A and B). Similarly, there was an overall trend of networks located at different positions along the human disturbance gradient having different species composition (tables S4, S5 and fig. S9C). Despite these effects, spatial distance alone accounted for the greatest proportion of deviance explained in species turnover across networks, followed by the shared contribution of spatial distance and ecoregion boundaries (fig. S10).

We found that plant-frugivore interaction dissimilarity increased significantly across ecoregions, biomes, and different levels of human disturbance, even after accounting for the effects of spatial distance, elevational differences, and sampling-related metrics (Table 1 and table S6). This provides strong support to the hypothesis that large-scale ecological boundaries mark spatially abrupt changes in plant-frugivore interactions (Figs. 1, 2 and fig. S11). Importantly, a great proportion of the deviance explained by biomes was shared with ecoregions (Fig. 3 and fig. S12), which suggests that changes in interaction dissimilarity across biome boundaries mostly reflect the variation occurring at a finer (ecoregion) scale. In fact, crossing an ecoregion boundary induced an average 7% increase in interaction dissimilarity, while crossing a biome boundary induced only an additional 0.2% change. As with species, networks located at opposite ends of the human disturbance continuum usually exhibited very different interactions (Fig. 4 and fig. S13).

**Fig. 1.**
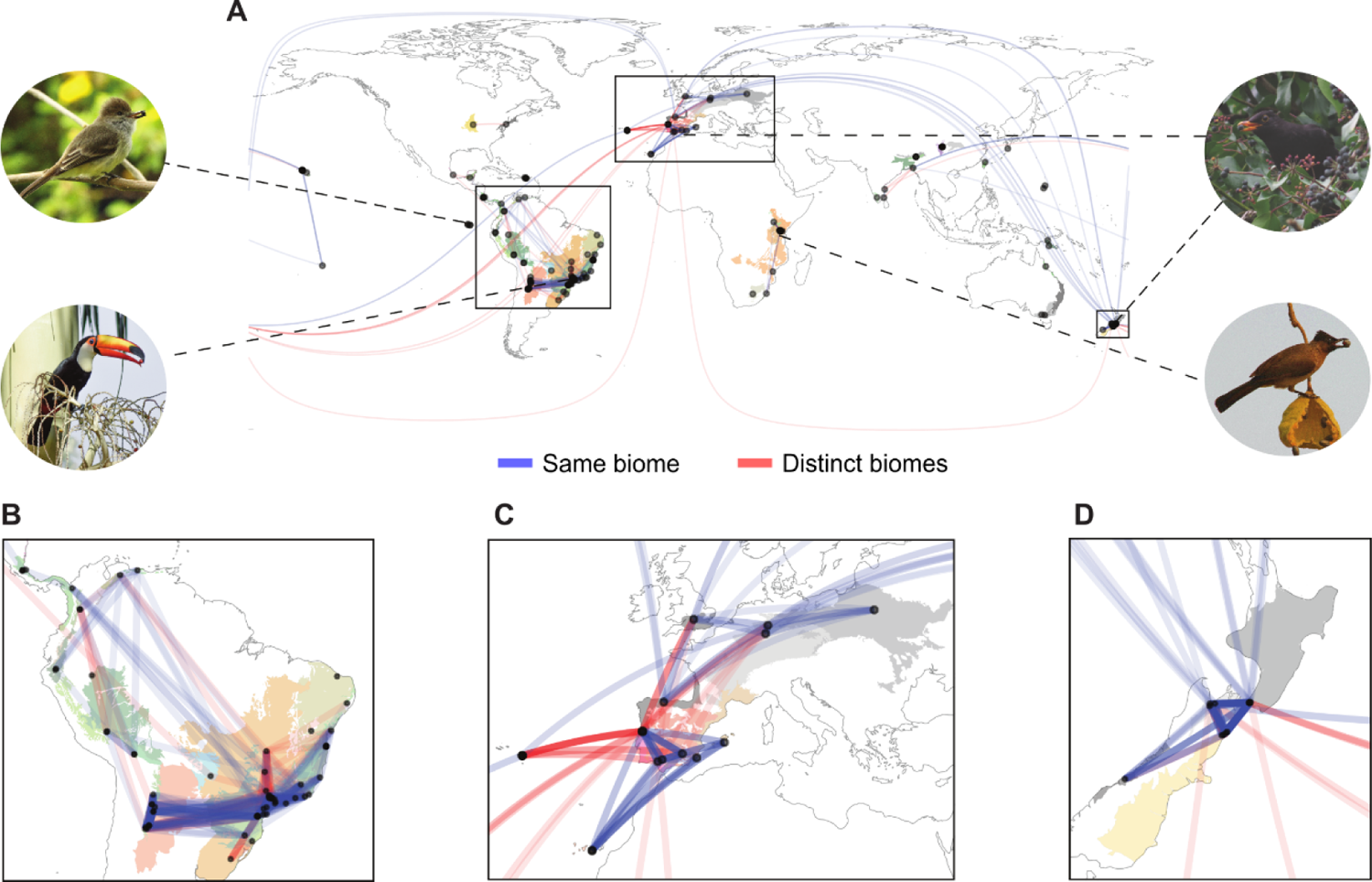
Plant-frugivore interactions shared among local networks, ecoregions and biomes. (**A**) World map with points representing the 196 local avian frugivory networks in our dataset. Colors of shaded areas represent the 67 ecoregions where networks were located, with similar colors indicating ecoregions that belong to the same biome. Lines represent the connections (shared interactions) plotted along the great circle distance between networks, with most of these connections occurring within (blue lines) rather than across (red lines) biomes. Stronger colour tones of lines indicate higher similarity of interactions (1-β_WN_) between networks. Connections across continents were mostly attributed to introduced species in one of these regions. Photos show some of the frugivorous birds present in our dataset. Inset maps depict three regions with many networks and connections (especially within biomes). (**B**) South America. (**C**) Europe. (**D**) Aotearoa New Zealand. [Photo credits: R. Heleno (top left and bottom right); R. B. Missano (bottom left); J. M. Costa (top right)].

**Fig. 2.**
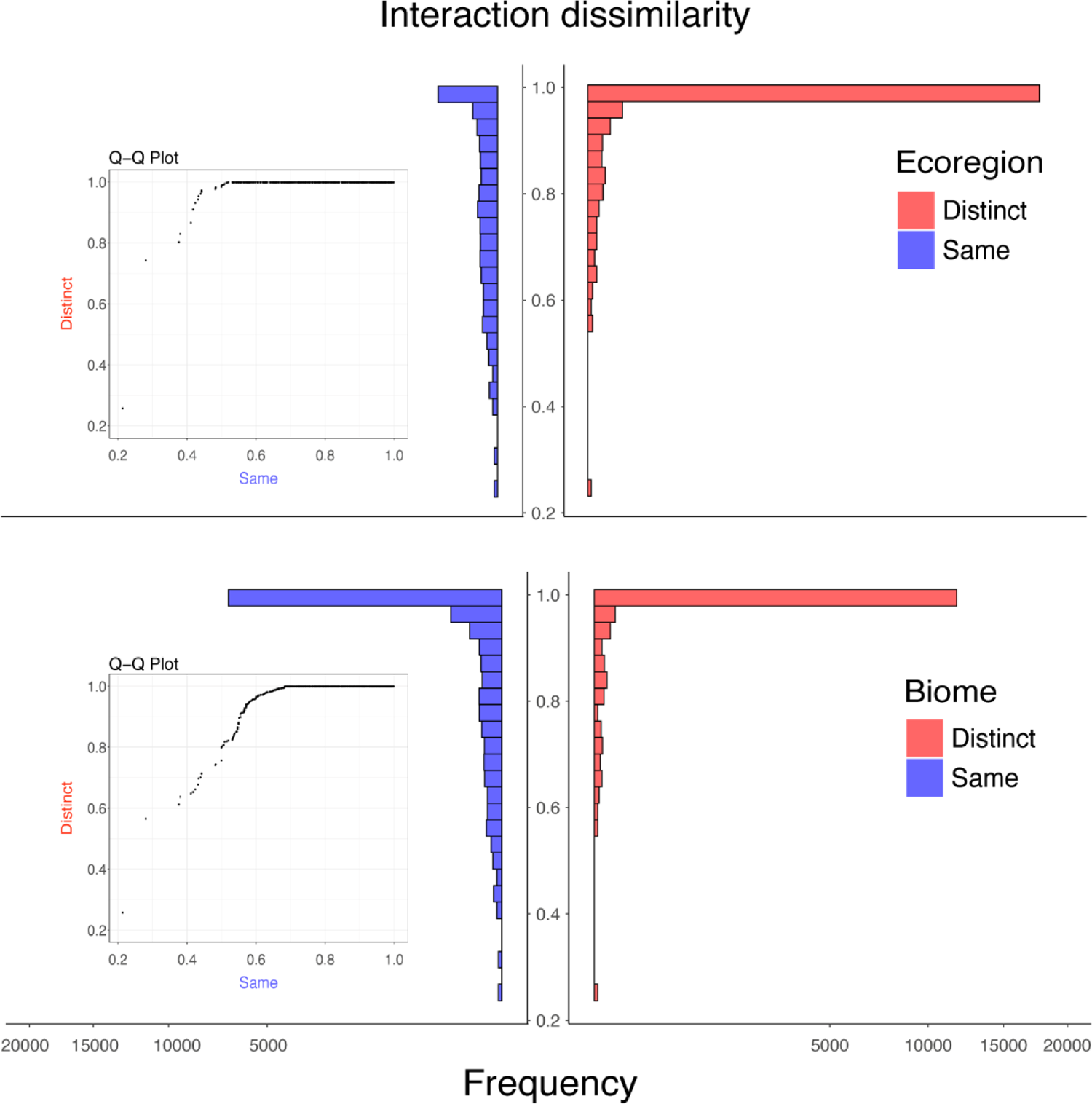
The effects of ecological boundaries on interaction dissimilarity. Histograms and inset quantile-quantile plots showing differences in the distributions of interaction dissimilarity values between pairs of networks located within (‘same’) and across (‘distinct’) ecoregions and biomes. The effects of ecoregion and biome boundaries were significant, even after controlling for the other predictor variables in the model. We square root transformed the x-axis scale to allow a better visualization of the distribution of data points (pairs of networks) with interaction dissimilarity values < 1.

**Fig. 3.**
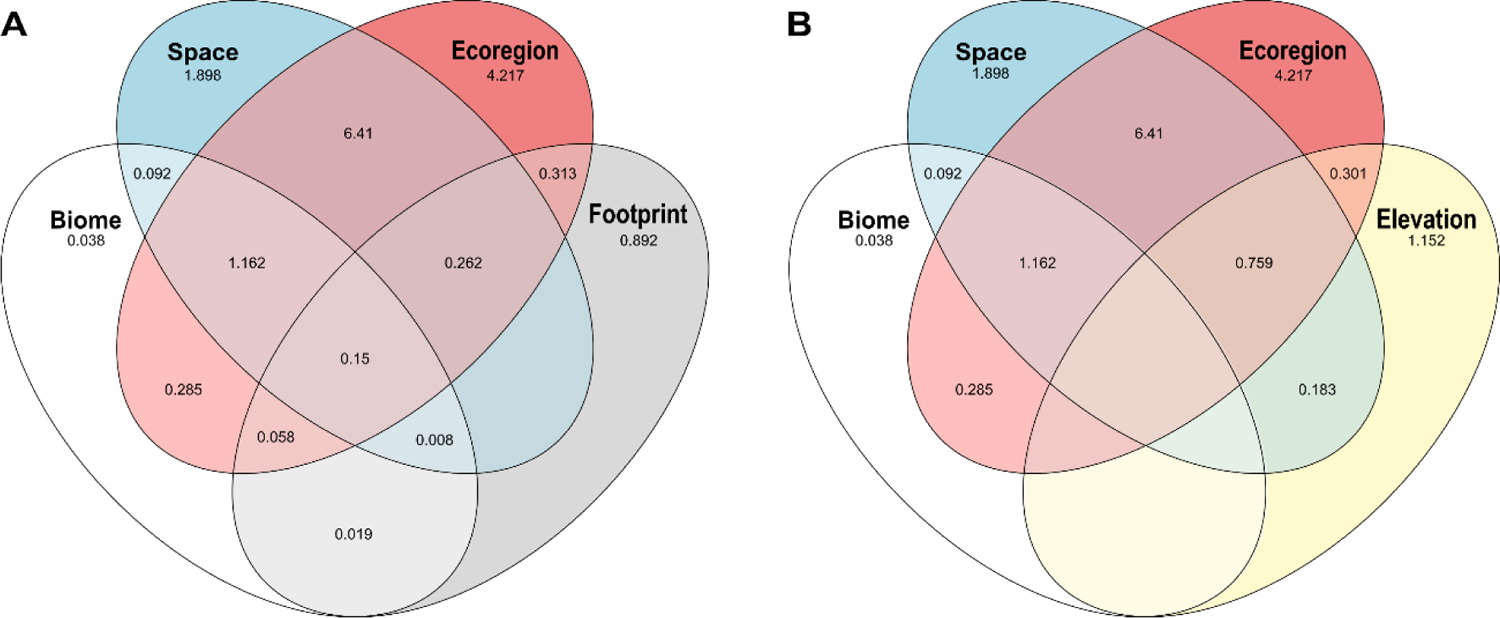
Venn diagrams showing the relative contributions (%) of our main predictor variables to explaining the variation in interaction dissimilarity, calculated using deviance partitioning. Overlapping areas represent deviance that is jointly explained by one or more predictor variables. (A) The relative contributions of ecoregions, biomes, spatial distance and human disturbance (i.e., footprint). In (B), we replace human footprint distance with elevational difference; we show these two separate diagrams for visualization purposes, but fig. S12 shows the effect of all our main predictor variables together. Note that we only plot our predictor variables of interest (i.e., not those used for controlling sampling effects). Terms that reduce explanatory power are not shown.

**Fig. 4.**
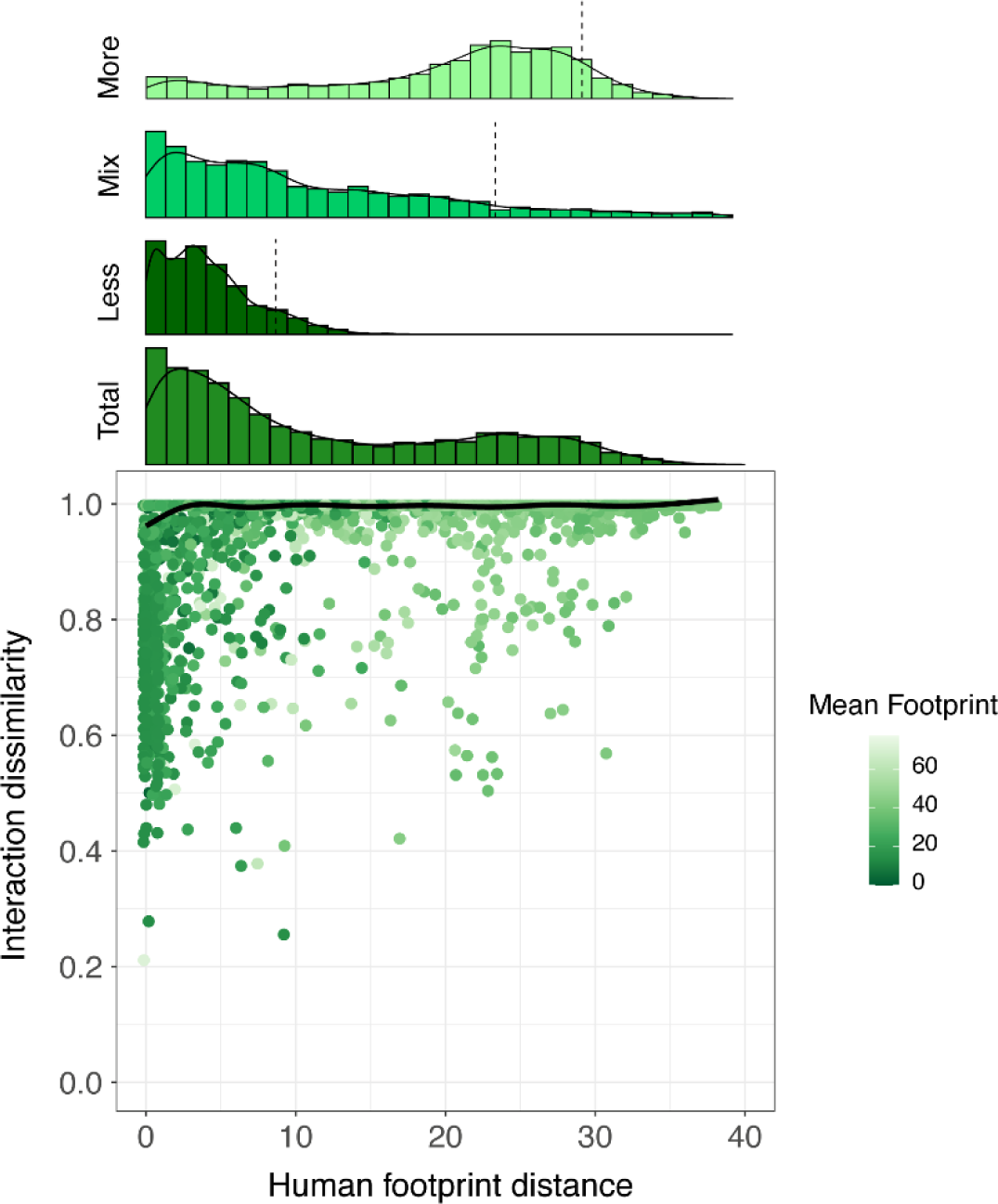
The effect of human disturbance gradients on interaction dissimilarity. The relationship between human disturbance (i.e., footprint) distance and interaction dissimilarity, with a fitted line obtained from a generalized additive model (GAM) with human footprint distance as the only predictor variable (fig. S13 shows the partial effects plot for the model including all predictors). Each data point (pair of networks) is coloured according to the mean of the human footprint values from the two networks. The histogram above the plot shows the distribution of data points across the human disturbance gradient. We further divided our data into three equal sized groups (top three histograms) based on their mean footprint values: ‘Less’ disturbed (low mean footprint), ‘Mix’ (medium mean footprint) and ‘More’ disturbed (high mean footprint). Dashed lines mark the 90^th^ percentile position in each histogram. Note that data points from less disturbed site pairs are skewed towards low values of human footprint distance, whereas pairs of more disturbed sites also had a larger average distance.

In addition to the importance of ecological boundaries and human disturbance gradients in driving plant-frugivore interaction dissimilarity, these effects were observed against a background of increasing interaction dissimilarity through space. Indeed, interaction dissimilarity increased sharply until a threshold distance of around 2,500 km between network sites, beyond which few networks shared any interactions and dissimilarity remained close to its peak (Fig. 5 and fig. S14). In the cases where spatially distant networks shared interactions, these typically involved species that had been introduced in at least one location. For instance, the interaction between the Blackbird *Turdus merula* and the Blackberry *Rubus fruticosus* was shared between networks located more than 18,000 km apart: while both species are native in Europe, they have been introduced to Aotearoa New Zealand.

**Fig. 5.**
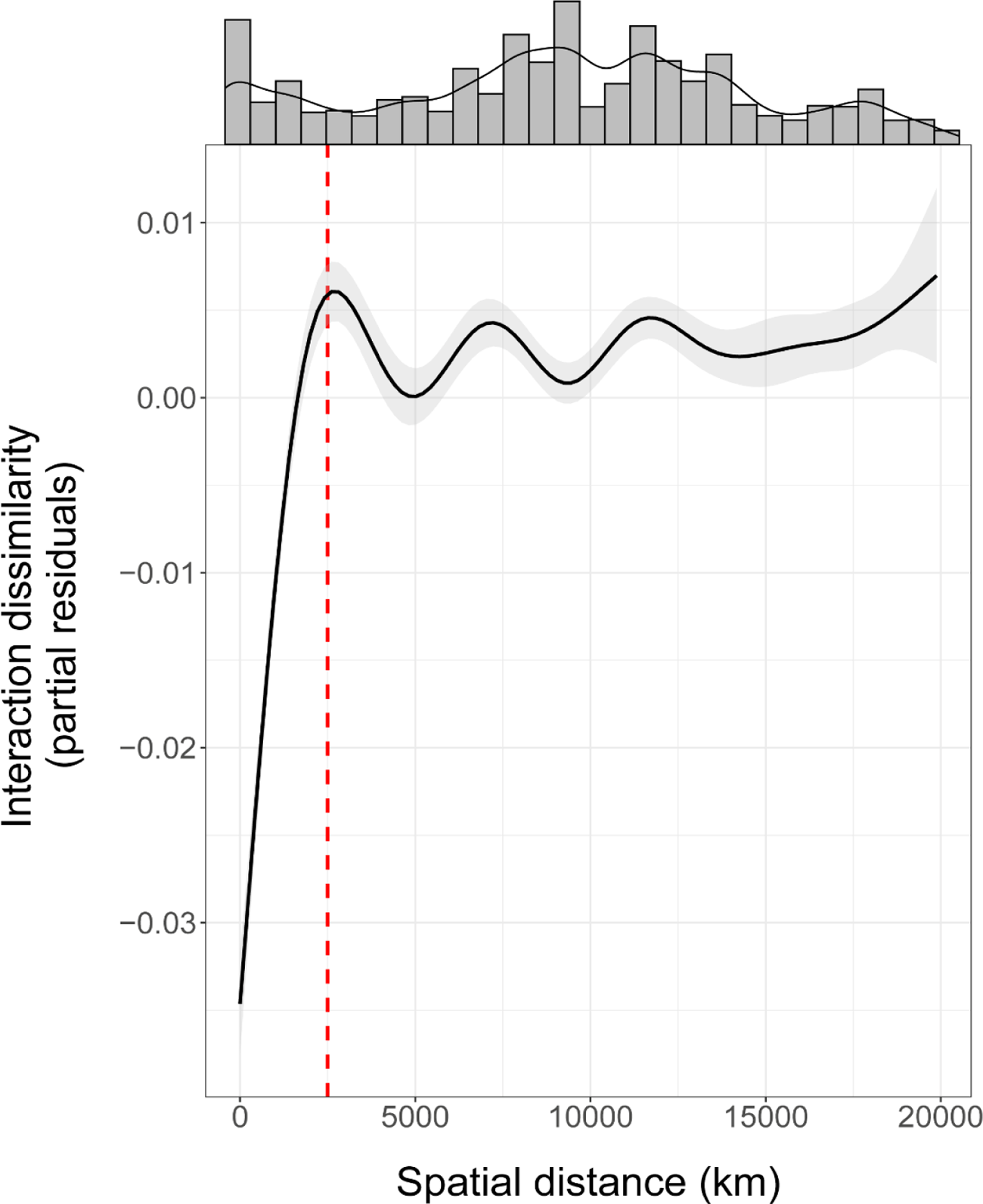
Partial effects plot of the relationship between spatial distance and interaction dissimilarity. Here, we show the fit (solid line) of a generalized additive model (GAM) with interaction dissimilarity as response variable and all our predictor variables included. Thus, this plot shows the effect of spatial distance on interaction dissimilarity, while controlling for the effect of the other predictors in the model. The gray area represents the 95% confidence interval of the fitted GAM. The histogram above the plot shows the distribution of data points across the spatial gradient. Note the sharp increase in interaction dissimilarity until a threshold distance of around 2,500 km (dotted red line), beyond which few networks shared interactions (also see fig. S14).

The shared effect of ecoregion boundaries and spatial distance explained the greatest proportion of the variability in plant-frugivore interaction dissimilarity, followed by the unique contributions of these two variables (Fig. 3). This indicates that gradual increases in interaction dissimilarity over space are made significantly steeper when crossing ecoregion boundaries; this combined effect better describes the variation in plant-frugivore interactions at large spatial scales than any of the other analysed variables. Despite significant turnover in species and interaction composition, structural dissimilarity of frugivory networks did not change consistently across large-scale ecological boundaries and human disturbance gradients, being only affected by spatial and sampling intensity distances (tables S7 and S8). All the above findings held true when evaluating both the binary and quantitative versions of ecoregion and biome distance matrices (Table 1 and tables S4 to S8). Furthermore, all our main results were robust to different processes of assigning uniqueness to problematic species (i.e., species without a valid epithet) (tables S9 to S33). Finally, most reported patterns were robust to the removal of individual studies from the dataset (figs. S15 and S16; tables S34 and S35).

Our results support the hypothesis that large-scale ecological boundaries drive abrupt changes in species and interaction composition of avian frugivory networks. Specifically, on top of the gradual effect of spatial distance on interaction dissimilarity (whereby networks > 2,500 km apart had very few interactions in common), transitions across ecoregions and biomes promoted divergence in species interactions. These results show that ecoregions and biomes, classically defined based on environmental conditions and species occurrences (*1, 3, 4*), also carry a signature within biotic interactions. This means that species biogeography is matched by a higher-order biogeography of interactions. In parallel, human disturbance gradients promoted shifts in species and interaction composition, which might be partly attributed to the filtering of sensitive species and their interactions from disturbed sites (*17, 27*). In fact, while networks from natural ecosystems usually contain interactions between native species, which better reflect natural biogeographic patterns (*10*) and are more susceptible to human disturbances (*27*), interactions from high-disturbance regions are generally performed by generalist and introduced species (*17, 27, 28*). Nevertheless, we found that the structure of avian frugivory networks was relatively consistent across large-scale environmental gradients. Similar results have been reported at smaller scales (*28*), indicating that assembly rules may generate common structural patterns in plant-frugivore networks (*29*) despite the shifts in species and interaction composition that usually accompany environmental changes (*13*).

Because most of the variation in interaction dissimilarity across biome borders can be explained by ecoregion boundaries, preserving the distinctness of ecoregions (*3, 4*) will likely contribute to maintaining the natural barriers that limit the spread of disturbances across the global network of frugivory. Unfortunately, the unique assemblages that comprise ecoregions have been increasingly threatened by global changes (*4, 5*). Biotic homogenization, in particular, has contributed to blurring biogeographical signatures (*10, 15*) and the effect of spatial processes on interaction dissimilarity (*10*). This notion is reinforced by the fact that all long-distance (>10,000 km) connections (shared interactions) between local networks of frugivory involved at least one region where novel interactions performed by introduced species have largely replaced those performed by native species, such as Aotearoa New Zealand and Hawai’i (*28, 30*). Interestingly, these long-distance connections tend to occur more frequently within than across biomes, despite a greater proportion of network comparisons being cross-biome (fig. S17). This indicates that biomes may represent meaningful boundaries not only for species, but also for novel interactions resulting from species introductions around the world (*10*). Taken together, these results suggest that disturbances in local networks of frugivory are much less likely to impact networks from distant sites or elevations, especially if the networks are located within distinct ecoregions and biomes.

Although species turnover and interaction dissimilarity responded to similar ecological drivers, species might interact differently across environmental gradients not only because of changes in species composition, but also because of partner switching associated with shifts in species abundance (i.e., the probability of random encounters), foraging behaviour and co-evolutionary patterns (*13*). To evaluate whether interaction rewiring [i.e., the extent to which shared species interact differently (*25*)] increases across large-scale environmental gradients, we used data limited to pairs of networks sharing plant and bird species (n = 1,314) (*23*). Of these, 93% had some degree of interaction rewiring, while around 30% did not have any interaction in common. We found that interaction rewiring increased significantly across human disturbance, spatial, and elevational gradients (table S36), partially explaining why interactions tend to turn over faster than species at large spatial scales (figs. S9D and S14C). Indeed, networks shared considerably more species than interactions (Fig. 1 and fig. S18), reinforcing previous findings that plant and bird species are flexible and tend to switch among their potential partners, even when networks have similar species composition (*28*). Surprisingly, we did not find an effect of ecoregion boundaries on interaction rewiring, probably because of their collinearity with our other predictor variables (tables S36 and S37).

As with other studies [e.g., (*10, 31*)], our data were not evenly spread across the globe, which likely affected the observed patterns. For instance, around 59% of our networks were located within a single tropical biome (fig. S2). Because ecoregions tend to be more distinct in tropical than in temperate zones (*32*), the greater number of networks from tropical ecosystems [which also possess most of the world’s ecoregions (*4*)] may have contributed to the strong observed effect of ecoregion boundaries on interaction dissimilarity. Nevertheless, both species richness and the proportion of frugivorous birds reach their peaks in the Tropics (*33*), suggesting that the distribution of networks in our dataset partially mirrors the global distribution of avian frugivory. Importantly, the extent to which our results apply for other frugivorous taxa (such as mammals and reptiles) and interaction types remains to be investigated. Previous findings, however, indicate that less mobile taxa tend to show a stronger adherence to ecological boundaries (*32*), a pattern that is likely to be reflected in species interactions.

This work provides evidence that ecological boundaries and human disturbance gradients delineate the large-scale distribution of species and their interactions. Nevertheless, network structure remained consistent across environmental gradients, suggesting that the ecological processes underlying the architecture of frugivory networks may be independent of species and interaction composition. By demonstrating the validity of the ecoregion-based approach (*4, 5*) for species interactions, our results have important implications for maintaining the world’s biodiversity of interactions and the myriad ecological functions they provide.

## Acknowledgments

We thank all the researchers in Tylianakis lab for their insightful comments on this work.

## Author contributions

Conceptualization: LPM, JMT Methodology: LPM, JMT, DBS Investigation: LPM, JMT, DBS Visualization: LPM, JMT Funding acquisition: LPM, JMT Project administration: JMT Supervision: JMT, DBS Writing – original draft: LPM, JMT Writing – review & editing: All authors.

## Data and materials availability

The data and code used in this study will be available for download at dryad (*34*).

## Supplementary Materials

Materials and Methods Figs. S1 to S18

Tables S1 to S37 References (35–171)

