## Supplementary Materials for "Global and regional ecological boundaries drive abrupt changes in avian frugivory interactions"

**This PDF file includes:**

Materials and Methods
Figs. S1 to S18
Tables S1 to S37

### Materials and Methods

Dataset acquisition

Plant-frugivore network data were obtained through different online sources and publications (Table S1). Only networks that met the following criteria were retrieved: i) the network contains quantitative data (a measure of interaction frequency) from a location, pooling through time if necessary; ii) the network includes avian frugivores (all other taxa were removed from analyses); iii) the network (after removal of non-avian frugivores) contains more than two species in each trophic level, and iv) network sampling was not taxonomically restricted, that is, sampling was not focused on a specific taxonomic group, such as a given plant or bird family. Note, however, that authors often select focal plants or frugivorous birds to be sampled, but this was not considered as a taxonomic restriction if plants and frugivores were not selected based on their taxonomy (e.g., focal plants were selected based on the availability of fruits at the time of sampling, or focal birds were selected based on previous studies of bird diet in the study site). The first source for network data was the Web of Life database (*35*), which contains 33 georeferenced plant-frugivore networks from 28 published studies, of which 12 networks met our criteria.

We also accessed the Scopus database on 04 May 2020 using the following keyword combination: (“plant-frugivore*” OR “plant-bird*” OR “frugivorous bird*” OR “avian frugivore*” OR “seed dispers*”) AND (“network*” OR “web*”) to search for papers that include data on avian frugivory networks. The search returned a total of 532 studies, from which 62 networks that met the above criteria were retrieved. We also contacted authors to obtain plant-frugivore networks that were not publicly available, which provided us a further 110 networks. We complemented our dataset by checking the references from a recently published study (*10*). Finally, 196 quantitative avian frugivory networks were used in our analyses (Fig. S2).

Standardizing the taxonomy

Considering the variety of authors and studies in our dataset, which identified plants and birds with differing resolution, it was necessary to reduce the taxonomic uncertainty in a uniform way. For this, we extracted the frugivore and plant species lists from all networks and performed a series of filters in order to remove non-existent species names (e.g., morphospecies labels) and standardize synonymous names according to reference databases.

*Frugivore species*

To account for spelling errors, we checked the matching of frugivore species names in our database to those from several taxonomic sources using the Global Names Resolver (GNR) (*36*). We accessed this database using the function *gnr_resolve* from the ‘taxize’ package in R (*37*) (Fig. S3; step 1). This function provides a matching score and the name from any of GNR’s sources that most closely matches each name in our species list. Matching is determined by a combination of checking for exact matches against the names in the data sources and fuzzy matching (of canonical forms or parts of the names) using the TaxaMatch algorithm (*38*). Because we were only interested in birds, we used the function *classification* from the same package to retrieve the taxonomic hierarchy and remove non-avian species, using the National Center for Biotechnology Information (NCBI) (*39*) as the reference database (Fig. S3; step 2). For those species classified as birds, we used the function *gnr_resolve* one more time using BirdLife International (*40*) as the reference database (Fig. S3; step 3). We used data from the Integrated Taxonomic Information System (ITIS) (*41*) and the *synonyms* function from the ‘taxize’ package (*37*) to obtain the synonyms of the species cross-checked with BirdLife International, as well as of those that were not found in the BirdLife database but were previously classified as birds (Fig. S3; step 4). We did this because, while obsolete bird species names usually did not have a match in BirdLife, one of its synonyms could: e.g., the black-fronted piping-guan was not found in the BirdLife database when its former scientific name, *Penelope jacutinga*, was entered; however, its currently accepted scientific name, *Pipile jacutinga*, was found as being one of the synonyms of *Penelope jacutinga*, and this synonym was revealed during step 4.

We also downloaded the Handbook of the Birds of the World (HBW) and BirdLife International (*42*) and automatically checked for matches of species names in our frugivore list with the names from the columns ‘scientific name’ and ‘synonym’ of the HBW-BirdLife spreadsheet (Fig. S3; step 5). By doing this, we were able to retrieve all the scientific names associated with the matched name in HBW-Birdlife. We used a fuzzy matching algorithm based on the Levenshtein distance between two strings to search for other possible names on the HBW-BirdLife spreadsheet for the species without good matches in any of the GNR’s sources or BirdLife International, as well as for those species that were not found in the ITIS database (Fig. S3; step 6). On some occasions, even this fuzzy matching algorithm could not find matches for a species name, which usually occurred when the genus name was incorrect or obsolete (note that in the vast majority of cases obsolete scientific names were fixed during steps 4 and 5, but some obsolete names were not present in either the ITIS or HBW-BirdLife databases). For those species, we automatically searched for their epithet names in the columns ‘scientific name’ and ‘synonym’ of HBW-BirdLife and retrieved only those that had one single match in each column (Fig. S3; step 7). The reason for restraining our search for those with one single match is because some epithet names are common and do not necessarily represent the same species. However, even this restriction is not a guarantee that the species with a given epithet in our list is the same species with the epithet in HBW-Birdlife, since a misspelled epithet name may coincidentally match the epithet of other species. Thus, we checked manually the taxonomy of all species corrected using this method (n = 17 species). We did this by searching for both the original species name (before the data cleaning process) and the matched name in the Avibase (*43*) and BirdLife (*40*) databases. By applying this series of filters, we were able to correct and validate the names and synonyms of 1,019 bird species. For the remaining 16 species, we checked the taxonomy manually by inspecting the same databases as in the previous step.

Finally, we generated a list object in R (*44*) in which element names correspond to scientific names accepted by either BirdLife International - obtained using the *gnr_resolve* function from the ‘taxize’ package (*37*) in 28/07/2020 - or HBW-BirdLife (*42*), while strings within elements correspond to all their synonyms and original species names. We used this list to standardize the taxonomy of the bird species in our local networks, so that synonyms would not be treated as different species. All species that were removed during the cleaning process (non-bird species and those without genus and/or species names, such as Undefined sp. and *Turdus* sp.) were removed from our local networks and further analyses (n = 82 species). Around 86% of frugivore species remained per network after the data cleaning (including the further removal of frugivore species for which all interaction partners were lost during the data cleaning process). Figure S3 shows a summary of the steps of the frugivore data cleaning.

*Plant species*

We checked the matching of plant species names with several taxonomic sources from the Global Names Resolver (GNR) (*36*) using the function *gnr_resolve* from the 'taxize' package (*37*) (Fig. S4; step 1), as with birds above. For those species without matches in any of GNR's sources, we applied a fuzzy matching algorithm based on the Levenshtein distance between two strings to compare these species' names with the matched names from GNR (Fig. S4; step 2). We did this because some of the species' names without matches in our step 1 were misspelled names of plant species already included in our dataset but not found by the *gnr_resolve* function. After this process, we relied on the *gnr_resolve* function one more time to compare the list of matched names from GNR with the list from the International Plant Names Index (IPNI) (*45*) (Fig. S4; step 3). The reason for using *gnr_resolve* twice is because we first wanted to make sure that the species had a match with at least one of the taxonomic sources from GNR (i.e., confirm that it is a scientific name) and then check whether the matched name represents a scientific name accepted by IPNI. By doing this, we were able to evaluate which species had high matching scores during our first step but not during the third, indicating that they are not internationally accepted scientific names.

We used data from the Tropicos database (*46*) to obtain the synonyms of the plant species that had been cross-checked with IPNI. We also relied on the IPlant collaborative database (*47*) to complement the synonyms list and retrieve the most recent accepted names of the species (Fig. S4; step 4). Using this series of filters, we were able to correct and validate the names and synonyms of 1,562 plant species. Finally, we generated a list object in R (*44*) in which element names correspond to accepted scientific names of species (cross-checked with the IPNI database on 15/09/2020) and strings within elements correspond to all their synonyms and original species names (before the data cleaning process). We used this list to standardize the taxonomy of the plant species in our local networks, as we did for birds.

Because our plant list contained cases in which two (or more) accepted species shared a synonym within their elements (n = 121), we had to deal with the standardization of these names. We did this by attributing the same name for all the occurrences of the species sharing a synonym only if the shared name was already present in our dataset. For example, *Cecropia digitata* is one of the synonyms of *C. angustifolia*, *C. obtusifolia* and *C. pachystachya* (and is therefore within the elements of these three species), but *C. digitata* was not present in any of the networks in our dataset, such that we could maintain the names C. *angustifolia, C. obtusifolia* and *C. pachystachya* in our local networks. We did this because shared synonyms that were not present in our dataset usually represented obsolete species that are no longer accepted. Alternatively, for the cases in which the shared synonym was present in our dataset (n = 37), we attributed the same name in the local networks for all the species that shared that given name. We adopted this conservative approach because, in this case, shared synonyms were usually species that were described multiple times by different authors, or species with several subspecies and varieties (note, however, that authors rarely include this level of taxonomic information on networks). Therefore, the shared name could potentially be any of the species that possess it as one of its synonyms.

Considering the high number of species (n = 184) with a valid genus name but without a valid epithet name (as indicated by the absence of matches in our steps 1 and 3, or by the low matching scores to any of the GNR's sources), as well as unresolved species names without good matches in the IPNI database (hereafter, *problematic species*) in our plant species list, we added two steps to evaluate whether such problematic species could be considered as a separate species from the other species in our dataset. For example, a species without an epithet (e.g., labelled in a study as '*Miconia* sp.') could still be treated as a distinct species in the analysis, provided we could be certain that it was not the same as another congeneric (*Miconia*) species, with or without epithet, in our dataset. Similarly, an unresolved species name that is not internationally accepted could only be considered as a distinct species in our analysis if we could disentangle it from its congeneric species in the dataset. Importantly, we did not perform these additional steps for birds because there were only a few cases of birds with valid genus but invalid epithet names.

To determine whether problematic species could be treated as a distinct species for analysis, we evaluated whether the distribution of any of the congeners of problematic species in our dataset overlapped with the location of the problematic species, such that we cannot be confident that the problematic species is not simply another occurrence of one or more of its congeners already in the dataset. For this process, we used the coordinates of the networks in which each problematic species occurred and generated buffer zones (diameter = 500 km) around these network locations. Considering that the size of the buffer zones could potentially affect our results, we also conducted the analysis using buffer zone sizes of 100 km and 1000 km (note, however, that our results still hold independently of the buffer zone size used; Tables S9 to S33). We collected occurrence data for all other species in the same genus in our dataset to evaluate whether the occurrence points of any of these congeneric species overlapped with the buffer zone of the problematic species (Fig. S4; step 5). For collecting occurrence points, we used data from the Global Biodiversity Information Facility (GBIF) (*48*) and applied a series of filters (for details, see the 'Occurrence data' section below). If the occurrence points of at least one of the congeneric species overlapped with the buffer zone of a given problematic species, we assumed that this problematic species could not be considered, with confidence, as a unique species in our dataset. Conversely, if none of the occurrence points of congeneric species overlapped with the buffer zone of the problematic species, we treated this problematic species as a separate species (Fig. S5), provided that there were no other problematic species (without valid epithets) in the same genus from other studies in the dataset.

Alternatively, if a genus contained more than one problematic species in the same study (e.g., *Miconia* sp.1, *Miconia* sp.2), we assumed that the authors distinguished the congeners within the study. For the cases in which a problematic species occurred in a single study and was the only species belonging to that genus in our dataset, the original name of the species was maintained in the local network. However, if there were problematic species from the same genus in different studies, we needed to ascertain whether they could potentially be the same species. Our approach for dealing with this issue was to determine all the possible species that a problematic species could be in each location, and then compare the lists of possible species in each location to identify any overlap. To do this, we first generated buffer zones (as in step 5) for each network location in which these problematic species occurred and obtained occurrence data from GBIF for all known species belonging to that genus (see the 'Occurrence data' section). We then checked whether there were congeneric species with occurrence points within the buffer zones of two (or more) problematic species belonging to the same genus (Fig. S4; step 6). If yes, we could not consider that these problematic species were different from each other. Rather, in this case there was a chance that the problematic species were the same species whose occurrence points overlapped the buffer zones of both network locations (Fig. S6). On the other hand, if there were no species whose distribution overlapped the buffer zones of both network locations, these problematic species could be considered as being distinct species in the dataset.

All species that were removed during the data cleaning process (i.e., the problematic species without a valid genus name, such as Rubiaceae sp. or Undefined sp.) were also removed from our local networks and further analyses (n = 166 species). Problematic species that could not be disentangled from resolved species or other problematic species in the dataset were named according to three distinct scenarios (for details, see the 'Alternative scenarios' section). Around 89% percent of plant species remained per network after the data cleaning (including the further removal of plant species for which all interaction partners were lost during the data cleaning process). Figure S4 shows a summary of the steps of the plant data cleaning.

*Occurrence data*

We retrieved occurrence data from the Global Biodiversity Information Facility (GBIF) (*48*) using the function *occ_search* from the R package 'rgbif' (*49*). For each species, we only requested occurrence data for observations for which coordinate points were available and no geospatial issues were detected, as determined by GBIF's record interpretation. We also followed a recent study (*50*) and removed occurrence points with: (i) a coordinate uncertainty larger than 100 km (the size of our smallest buffer zone); (ii) those for which the collection date was before 1945, as older occurrence points are usually not properly geo-referenced (*51*); (iii) those in which the number of counts associated with the occurrence point was zero; and (iv) those in which the 'basis of record' was not an observation or a preserved specimen.

In addition, we used the function *clean_coordinates* from the R package 'CoordinateCleaner' (*52*) and land mass and country data (with a 1:10m scale) from Natural Earth (*53*) to remove occurrence points for which the coordinates: (v) fell within the ocean or outside the borders of the country where they were recorded, both of which indicate data-entry errors, (vi) were located around the country capital or the centroid of the country, indicating imprecise geo-referencing based on inadequate sampling site descriptions, (vii) both latitude and longitude were zero or had equal values, indicating failed geo-referencing, and (viii) were located around a biodiversity institution, suggesting that records might represent specimens that were erroneously geo-referenced to museums, herbaria or universities instead of their sampling localities (*52*). After applying this series of filters, 913,1777 occurrence points were retrieved for 623 plant species in our dataset. These occurrence points were used for disentangling 'problematic' species during step 5 of the plant species cleaning process (Fig. S5).

Because the next step required us to retrieve occurrence data for all known species belonging to a given genus, we used the function *name_lookup* from the R package 'rgbif' (*49*) to search for all accepted species names associated with the genus name. We used the same set of filters previously described to obtain the occurrence points for each species during the step 6 of the plant species cleaning process (Fig. S6). In the end, 1,988,540 occurrence points were retrieved for 4,793 plant species.

*Alternative scenarios*

We used three distinct scenarios for attributing names for problematic plant species that could not be considered as unique species in our dataset. In the first scenario, we removed from the local network any problematic species whose buffer zone was overlapped by the distribution of 'resolved' congeneric species in the dataset (step 5 of the plant species cleaning process). For example, if the buffer zone of the problematic species '*Miconia* sp.' was overlapped by other resolved *Miconia* species in the dataset, we removed the species *Miconia* sp. (and all of its interactions) from its local network. We adopted this strategy rather than considering that the problematic species and the resolved species that overlap its buffer zone are the same because such problematic species could potentially be any of the resolved species that overlap its buffer zone. This, in turn, made it impractical to attribute the name of the resolved species to the problematic species in cases where the buffer zone of the problematic species was overlapped by several resolved species. In addition, our first scenario considers all problematic species that could not be disentangled from each other (step 6 of the plant species cleaning process) as being the same species. For example, if two problematic species labelled as '*Coussapoa* sp.' in two separate local networks could not be disentangled because there are congeneric species simultaneously overlapping the buffer zones of both network locations (Fig. S6), we attributed the same name to these two problematic species.

Alternatively, our second scenario treats problematic species as being unique. Therefore, a unique name was given for the problematic species whose buffer zone was overlapped by 'resolved' congeneric species in the dataset. For instance, the problematic species '*Miconia* sp.' from the example above would receive a unique name in the second scenario instead of being removed from its local network. In this scenario, we also attributed unique names for problematic species that could not be disentangled from each other. For example, each of the two problematic *Coussapoa* species mentioned above would receive a unique name instead of sharing the same name.

Finally, the third scenario removes from the local networks all plant species that could not be considered as being unique species in the dataset, and is therefore our most conservative scenario (which was used for obtaining the results presented in the main text). Because these three different scenarios could affect our response variables, we repeated the analyses using the sets of networks from all scenarios. Note, however, that results remained qualitatively the same independently of the scenario used in the analyses (Tables S9 to S33).

<u>Generating the distance matrices</u>

We generated several distance matrices (N × N, where N is the number of local networks in our dataset) to be our predictor and response variables in the statistical models. Here, we detail each one of them:

**Ecoregion and biome distances:** We used the most updated map of ecoregions and biomes (*4*), which divides the globe into 846 terrestrial ecoregions nested within 14 biomes, to generate our ecoregion and biome distance matrices. Of these, 67 ecoregions and 11 biomes are represented in our dataset (Figs. S1 and S2). We constructed alternative versions of both the ecoregion and biome distance matrices. In the binary version, if two ecological networks were from localities within the same ecoregion/biome, a dissimilarity of zero was given to this pair of networks, whereas a dissimilarity of one was given to a pair of networks from distinct ecoregions/biomes (this is the same as calculating the Euclidean distance on a presence-absence matrix with networks in rows and ecoregion/biomes in columns).

In the quantitative version, we estimated the pairwise environmental dissimilarity between our ecoregions and biomes using six environmental variables recently demonstrated to be relevant in predicting ecoregion distinctness, namely mean annual temperature, temperature seasonality, mean annual rainfall, rainfall seasonality, slope and human footprint (*32*). We obtained climatic and elevation data from Worldclim (*54*) at a spatial resolution of 1-km$^2$. We transformed the elevation raster into a slope raster using the function *terrain* from the 'raster' package (*55*) in R (*44*). Human footprint is a metric that combines eight variables associated to human disturbances on the environment: the extent of built environments, crop land, pasture land, human population density, night-time lights, railways, roads and navigable waterways (*24*). The human footprint raster was downloaded at a resolution of 1-km$^2$ (*24*). Because human footprint data were not available for one of our ecoregions (Galápagos Islands xeric scrub), we estimated human footprint for this ecoregion by converting visually interpreted scores into the human footprint index. We did this by analysing satellite images of the region and following a visual score criterion (*24*). Given the previously demonstrated strong agreement between visual score and human footprint values (*24*), we fitted a linear model using the visual score and human footprint data from 676 validation plots located within the Deserts and xeric shrublands biome (*24*) - the biome in which the Galápagos Islands xeric scrub ecoregion is located - and estimated the human footprint values for our own visual scores using the *predict* function in R (*44*).

We used 1-km$^2$ resolution rasters and the *extract* function from the 'raster' package (*55*) to calculate the mean value of each of our six environmental variables for each ecoregion in our dataset. Because biomes are considerably larger than ecoregions (which makes obtaining environmental data for biomes more computationally expensive) we used a coarser spatial resolution of 5-km$^2$ for calculating the mean values of environmental variables for each biome. Since a 5-km$^2$ resolution raster was not available for human footprint, we transformed the 1-km$^2$ resolution raster into a 5-km$^2$ raster using the function *resample* from the same package.

We ran a Principal Component Analysis (PCA) on our scaled multivariate data matrix (where rows are ecoregions or biomes and columns are environmental variables), selected the scores of the four and three principal components, which represented 89.6% and 88.7% of the variance for ecoregions and biomes, respectively, and converted it into a distance matrix by calculating the Euclidean distance between pairs of ecoregions/biomes using the *vegdist* function from the 'vegan' package (*56*). Finally, we transformed the ecoregion/biome distance matrix into a N × N matrix where N is the number of local networks. In this matrix, cell values represent the pairwise environmental dissimilarity between the ecoregions/biomes where the networks are located. The main advantage of using this quantitative approach is that, instead of simply evaluating whether frugivory networks located in distinct ecoregions or biomes are different from each other in terms of network composition and structure (as in our binary approach), we were also able to account for how different ecoregions and biomes are from one another.

**Human disturbance (footprint) distance:** To generate our local human footprint distance matrix, we extracted human footprint data at a 1-km$^2$ spatial resolution (*24*) and calculated the mean human footprint values within a 5-km buffer zone around each network site. For the networks located within the Galápagos Islands xeric scrub ecoregion (n = 4), we estimated the human footprint index using the same method described in the previous section for ecoregion- or biome-scale human footprint. We then calculated the pairwise Euclidean distance between human footprint values from our network sites. Thus, low cell values in the local human footprint distance matrix indicate pairs of network sites with a similar level of human disturbance, while high values represent pairs of network sites with very different levels of human disturbance.

**Spatial distance:** The spatial distance matrix was generated using the Haversine (i.e., great circle) distance between all pairwise combinations of network coordinates. In this matrix, cell values represent the geographical distance between networks.

**Elevational difference:** We calculated the Euclidean distance between pairwise elevation values (estimated as meters above sea level) of network sites to generate our elevational difference matrix. Elevation values were obtained from the original sources when available or using Google Earth (*57*). In the elevational difference matrix, low cell values represent pairs of network sites within similar elevations, whereas high values represent pairs of network sites within very different elevations.

**Network sampling dissimilarity:** We used the metadata retrieved from each of our 196 local networks to generate our 'network sampling dissimilarity matrices', which aim to control statistically for differences in network sampling. There are many ways in which sampling effort could be quantified, so we began by calculating a variety of metrics, then narrowed our options by assessing which of these was most related to network metrics. We divided the sampling metrics into two categories: time span-related metrics (i.e., sampling hours and months) and empirical metrics of sampling completeness (i.e., sampling completeness and sampling intensity), which aim to account for how complete network sampling was in terms of species interactions.

We selected the quantitative sampling metrics to be included in our models based on (i) the fit of generalized linear models evaluating the relationship between number of sampling hours and sampling months of the study and network-level metrics (i.e., bird richness, plant richness and number of links), and (ii) how well time span-related metrics, sampling completeness and sampling intensity predicted the proportion of known interactions that were sampled in each local network (hereafter, ratio of interactions) for a subset of the data. This latter metric, defined as the ratio between the number of interactions in the local network and the number of known possible interactions in the region involving the species in the local network, captures raw sampling completeness. Therefore, 'ratio of interactions' estimates the proportion of interactions involving a given set of species in the region's meta-network that are found in the local network. To calculate this metric, we needed high-resolution information on the possible interactions, so we used a subset of 14 networks sampled in Aotearoa New Zealand, since there is an extensive compilation of frugivory events recorded for this country (*58*). After this process, we selected number of sampling hours, number of sampling months and sampling intensity to be included in our statistical models (Figs. S7, S8 and Table S2). We generated these distance matrices by calculating the Euclidean distance between metric values.

Similarly, we generated a Euclidean distance matrix for differences in sampling year between networks, which aims to account for long-term changes in the environment, species composition and network sampling methods. We obtained the sampling year of our local networks from the original sources and calculated the mean sampling year value for those networks sampled across multiple years.

Because sampling methods, such as sampling design, focus (i.e., focal taxa), interaction frequency type (i.e., how interaction frequency was measured) and coverage might also affect the observed plant-frugivore interactions (*59*), we combined these variables into a single distance matrix to estimate the overall differences in sampling methods between networks. Considering that most of these variables were categorical with multiple levels (Table S3), we generated our methods dissimilarity matrix by using a generalization of Gower's distance method (*60*), which allows the treatment of different types of variables when calculating distances. For this, we used the *dist.ktab* function from the 'ade4' package (*61*). We ran a Principal Coordinates Analysis (PCoA) on this distance matrix, selected the first four axes, which explained 81.2% of the variation in methods dissimilarity, and calculated the Euclidean distance between pairs of networks using the *vegdist* function from the 'vegan' package (*56*) in R (*44*).

**Network dissimilarity:** We generated three network dissimilarity matrices to be our response variables in the statistical models. In the first, cell values represent the pairwise dissimilarity in species composition between networks (beta-diversity of species; $\beta_S$) (*25*). Second, we measured interaction dissimilarity (beta-diversity of interactions; $\beta_{WN}$), which represents the pairwise dissimilarity in the identity of interactions between networks (*25*). Importantly, we did not include interaction rewiring ($\beta_{OS}$) in our main analyses because this metric can only be calculated for networks that share interaction partners (i.e., it estimates whether shared species interact differently) (*25*), which limited the number and the spatial distribution of networks available for analysis (but see the 'Rewiring analysis' section). Metrics were calculated using the *network_betadiversity* function from the 'betalink' package (*62*) in R (*44*).

Finally, we calculated a third dissimilarity matrix to capture overall differences in network structure. We recognise that there are many potential metrics of network structure, and that many of these are strongly correlated with one another (*63–65*). We therefore chose a range of metrics that captured the number of links, their relative weightings (including across trophic levels), and their arrangement among species, then combined these into a single distance matrix. Specifically, we quantified network structural dissimilarity using the following metrics: weighted connectance, weighted nestedness, interaction evenness, PDI and modularity.

Weighted connectance represents the number of links relative to the number of possible links, weighted by the frequency of each interaction (*66*), and is therefore a measure of network-level specialization (higher values of weighted connectance indicate lower specialization). Importantly, it has been suggested that connectance affects persistence in mutualistic systems (*64*). We measured nestedness (i.e., the pattern in which specialist species interact with proper subsets of the species that generalist species interact with) using the weighted version of nestedness based on overlap and decreasing fill (wNODF) (*67*). Nested structures are common in plant-frugivore networks (*29*) and have been considered to increase the number of coexisting species by minimizing interspecific competition (*68*). Interaction evenness is the Shannon's evenness index applied for species interactions and represents how evenly distributed the interactions are in the network (*14*, *69*). This metric has been previously demonstrated to decline with habitat modification as a consequence of some interactions being favoured over others in high-disturbance environments (*14*). PDI (Paired Difference Index) is a measure of species-level specialization on resources and a reliable indicator not only of specialization, but also of absolute generalism (*70*). In order to obtain a network-level PDI, we calculated the weighted mean PDI for each local network. Finally, we calculated modularity (i.e., the level of compartmentalization within networks) using the DIRTPLAwb+ algorithm (*71*). Modularity estimates the extent to which species within modules interact more with each other than with species from other modules (*72*), and it has been demonstrated to affect the persistence and resilience of mutualistic networks (*64*). All the selected network metrics are based on weighted interaction data, as these have been suggested to be less biased to sampling incompleteness (*73*) and to better reflect environmental changes (*14*). All network metrics were calculated using the 'bipartite' package (*74*) in R (*44*).

We ran a Principal Component Analysis (PCA) on our scaled multivariate data matrix (N × M where N is the number of local networks in our dataset and M is the number of network metrics), selected the scores of the three principal components, which represented 89.9% of the variance in network metrics, and converted it into a network structural dissimilarity matrix by calculating the Euclidean distance between networks. In this distance matrix, cell values represent differences in the overall architecture of networks (over all the network metrics calculated), and therefore provide a complementary approach for evaluating how species interaction patterns vary across large-scale environmental gradients.

Statistical analyses

We employed a combination of Generalized Additive Models (GAM) and Multiple Regression on Distance Matrices (MRM) (*26*) to evaluate the effect of each of our predictor distance matrices on our response matrix. Essentially, this analysis is equivalent to a GAM, but where the predictor and response variables are distance matrices and the non-independence of distances from each given local network is accounted for in the hypothesis testing by permuting the response matrix. This analysis allowed us to obtain F-values for each of the smooth terms (i.e., smooth functions of the predictor variables in our model) and test statistical significance at the level of individual variables. The binary versions of ecoregion and biome distance matrices (with two levels, 'same' or 'distinct') were treated as categorical variables in the models, and t-values were used for obtaining statistical significance. We fitted GAMs with thin plate regression splines (*75*) using the *gam* function from the 'mgcv' package (*76*) in R (*44*). Smoothing parameters were estimated using restricted maximum likelihood (REML) (*76*). Our GAM-based MRM models were calculated using a modified version of the *MRM* function from the 'ecodist' package (*77*), which allowed us to combine GAMs with the permutation approach from the original function. All the models were performed with 1,000 permutations.

We explored the unique and shared contributions of our predictor variables to network dissimilarity using deviance partitioning analyses. These were performed by fitting reduced models (i.e., GAMs where one or more predictor variables of interest were removed) using the same smoothing parameters as in the full model and comparing the explained deviance. We fixed smoothing parameters for comparisons in this way because these parameters tend to vary substantially (to compensate) if one of two correlated predictors is dropped from a GAM.

Assessing the influence of individual studies on the reported patterns

Because our dataset comprises 196 local frugivory networks obtained from 93 different studies, and some of these studies contained multiple networks, we needed to evaluate whether our results were strongly biased by individual studies. To do this, we followed a previous approach (*78*) and tested whether F-values of smooth terms and t-values of categorical variables (ecoregion and biome) changed significantly when jackknifing across studies. We did this by dropping one study from the dataset and re-fitting the models, and then repeating this same process for all the studies in our dataset.

We found a number of consistent patterns within the data (Figs. S15 and S16); however, some of the patterns we observed appear to be driven by individual studies with multiple networks, and hence are less representative. For instance, the study with the greatest number of networks in our dataset (study ID = 76), which contains 35 plant-frugivore networks sampled across an elevation gradient in Mt. Kilimanjaro, Tanzania (*79*), had an overall high influence on the results when compared to the other studies. By re-running our GAM-based MRM models after removing this study from our dataset, we found that the effect of biome boundaries on interaction dissimilarity is no longer significant, whereas the effects of ecoregion boundaries, human footprint distance, spatial distance and elevational differences remained consistent (Table S34). Nevertheless, all the results were qualitatively similar to those obtained for the entire dataset when using network structural dissimilarity as the response variable (Table S35).

Rewiring analysis

Interaction rewiring ($\beta_{OS}$) estimates the extent to which shared species interact differently (*25*). Because this metric can only be calculated for networks that share species from both trophic levels, we selected a subset of network pairs that shared plants and frugivorous birds (n = 1,314) to test whether interaction rewiring increases across large-scale environmental gradients. Importantly, since not all possible combinations of network pairs contained values of interaction rewiring (i.e., not all pairs of networks shared species), a pairwise distance matrix could not be generated for this metric. Thus, we were not able to use the same statistical approach used in our main analyses, which is based on distance matrices (see 'Statistical analyses' section). Instead, we performed a Generalized Additive Mixed Model (GAMM) using ecoregion, biome, spatial, elevational, and sampling-related distance metrics as fixed effects and network IDs as random effects (to account for the non-independence of distances) (Table S36). We also performed a reduced model with only ecoregion and biome distance metrics as predictor variables (Table S37). The binary version of ecoregion and biome distance metrics (with two levels, 'same' or 'distinct') were used as categorical variables in both models. Interaction rewiring [$\beta_{OS}$, following (*25*)] was calculated using the *network_betadiversity* function from the 'betalink' package (*62*) in R (*44*). Although it has been recently argued that this metric may overestimate the importance of rewiring for network dissimilarity (*80*), our main focus was not the partitioning of network dissimilarity into species turnover and rewiring components, but rather simply detecting whether the sub-web of shared species interacted differently. In this case, $\beta_{OS}$ [as developed by (*25*)] is an adequate and useful metric (*80*). We fitted our models using the *gamm4* function from the 'gamm4' package (*81*) in R (*44*). Smoothing parameters were estimated using restricted maximum likelihood (REML) (*76*).

A

B

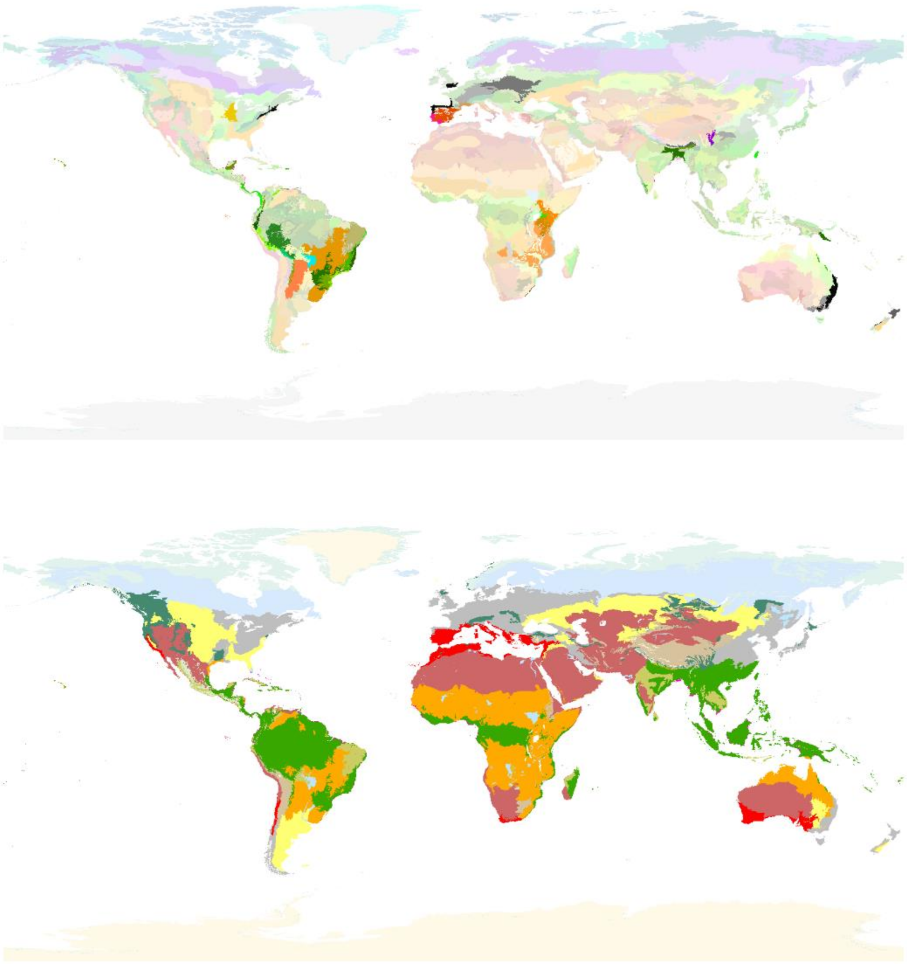

**Fig. S1. Maps of ecoregions and biomes of the world.** (**A**) Terrestrial ecoregions, with stronger colour tones indicating the 67 ecoregions represented in our dataset. (**B**) Global biomes, with stronger colour tones indicating the 11 biomes represented in our dataset. Boundaries were defined based on the most updated map of ecoregions and biomes (*4*).

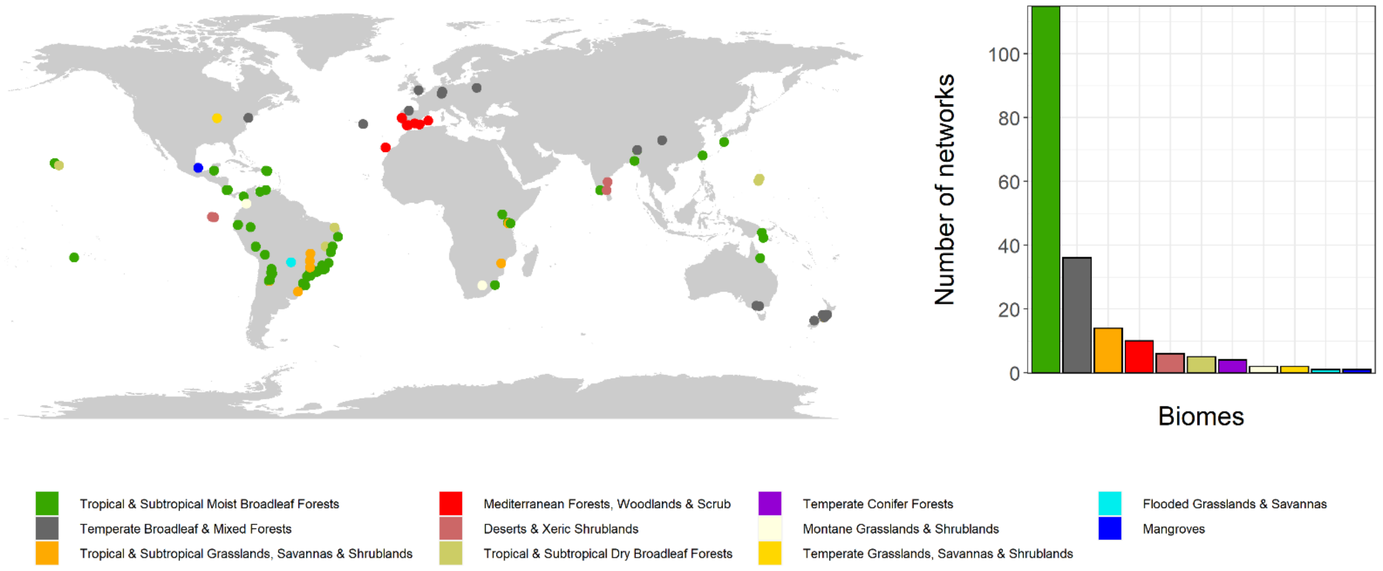


**Fig. S2. Geographic distribution of the 196 avian frugivory networks in our dataset.** Local networks were distributed across 11 biomes, with most of these being located within a single biome: the Tropical & Subtropical Moist Broadleaf Forests.

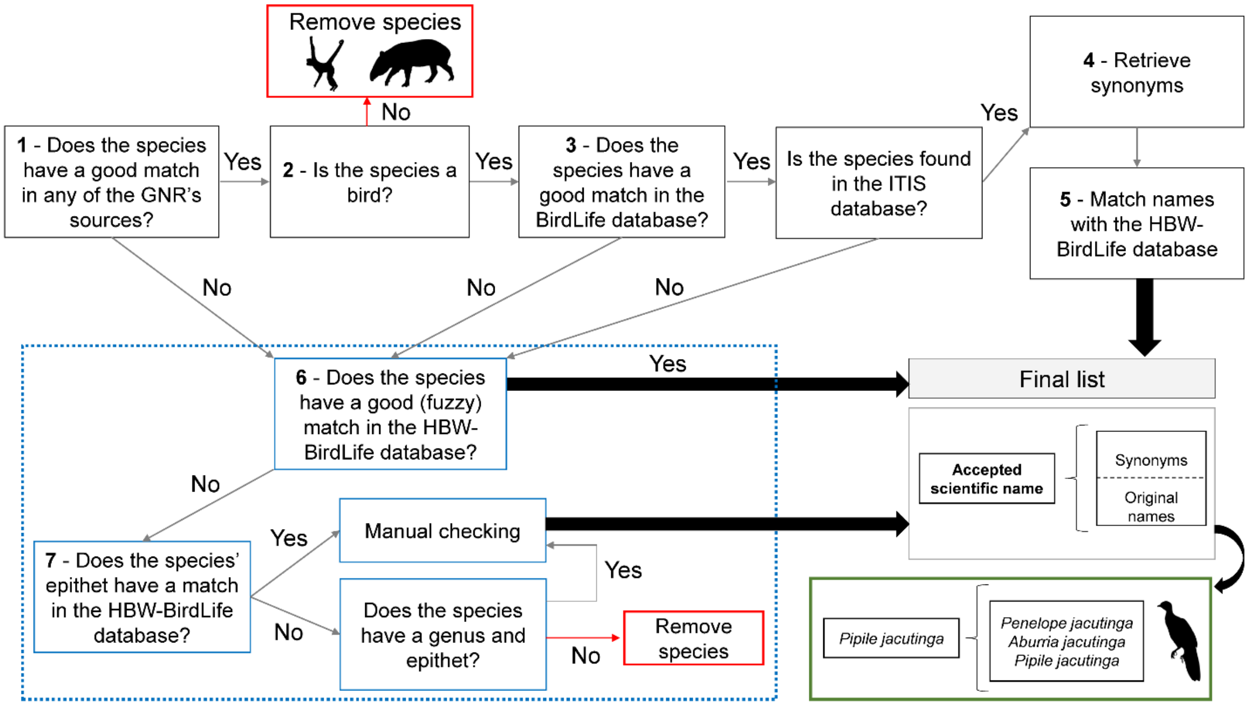


**Fig S3. An overview of the steps for cleaning and standardizing the frugivore species data.** Red boxes represent species that were removed from the analyses (non-avian species and species without epithet or genus names). The dashed box comprises the steps performed for the species without good matches in any of the Global Names Resolver (GNR) sources and in the BirdLife International database, or that were not found in the Integrated Taxonomic Information System (ITIS). The final list comprises elements whose names represent scientific names accepted either by the BirdLife International or by the Handbook of the Birds of the World and BirdLife International, and strings within elements comprise their synonymous and original names (before the cleaning process). For example, *Pipile jacutinga* (Cracidae) is the current accepted name of the black-fronted piping-guan, while its synonymous names include *Penelope jacutinga* and *Aburria jacutinga* (green box). All names (strings of synonyms) within elements (accepted names) were replaced by the element name in the local networks, such that a given species had the same name for all its occurrences in the entire database. Numbers inside boxes correspond to the steps of the frugivore data cleaning process (described above).

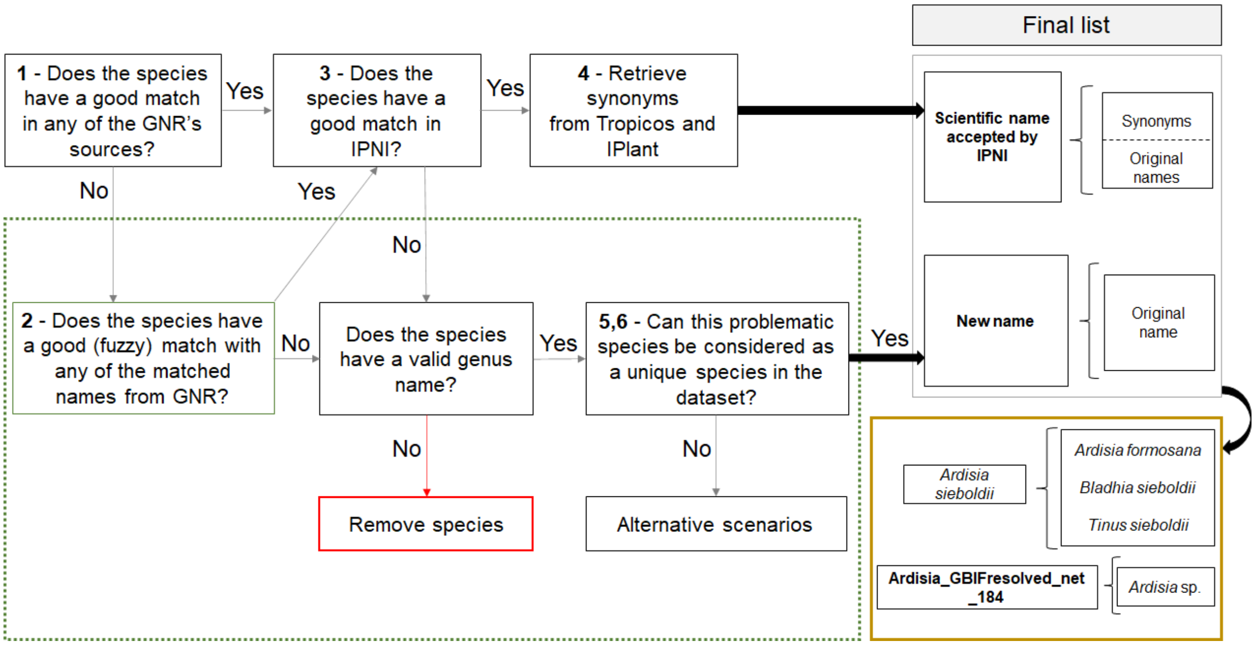


**Fig. S4. Overview of the steps for cleaning and standardizing the plant species data.** The red box represents species that were removed from the analyses (species without valid genus names). The dashed box comprises the steps performed for the species without good matches in any of the Global Names Resolver (GNR) sources and in the International Plant Names Index (IPNI) database (i.e., 'problematic species'). We performed two steps to determine if problematic species could be considered as being unique species in our dataset (see steps 5 and 6 of the plant species data cleaning process described in the text and visualized in Figs. S5 and S6). The final list comprises elements whose names represent scientific names cross-checked with the IPNI database and strings within elements comprise their synonymous and original names (before the cleaning process), or elements whose names represent new names given for problematic species that can be considered as unique species in our dataset, and the strings within elements comprise their original name. For example (yellow box), *Ardisia sieboldii* (Primulaceae) is a scientific name accepted by IPNI, while *A. formosana*, *Bladhia sieboldii* and *Tinus sieboldii* represent some of its synonymous names. Meanwhile, Ardisia_GBIFresolved_net_184 is the new name given for the problematic (but unique) species *Ardisia* sp., as revealed by the step 5 of the plant species data cleaning process. Note that, in the former case, all names (strings) within elements were replaced by the element name in the local networks, while in the latter case strings within elements were replaced by the element name only in the network where the problematic species was observed (in this example, network 184). Numbers inside boxes correspond to the steps of the plant data cleaning process. See the 'Alternative scenarios' section for details on how we attributed names for plant species that could not be considered as unique species in our dataset.

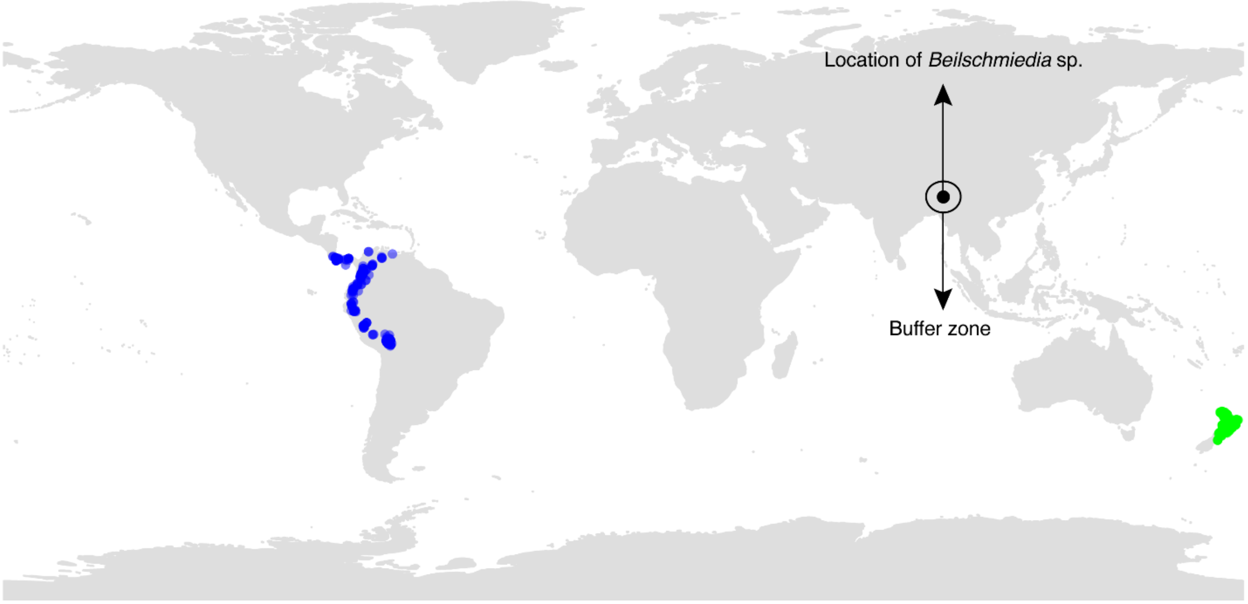


**Fig. S5. Graphical example (for an unresolved *Beilschmiedia* species) of step 5 of the plant species cleaning process**. Coloured points indicate the distribution of *Beilschmiedia* species already contained within our dataset, and these are compared with the occurrence location of a 'problematic species' (a species with genus name only). The distributions of both *Beilschmiedia tawa* (green dots) and *Beilschmiedia tovarensis* (blue dots) do not overlap with the buffer zone of the problematic species *Beilschmiedia* sp., such that *Beilschmiedia* sp. can be considered as a separate species in our dataset.

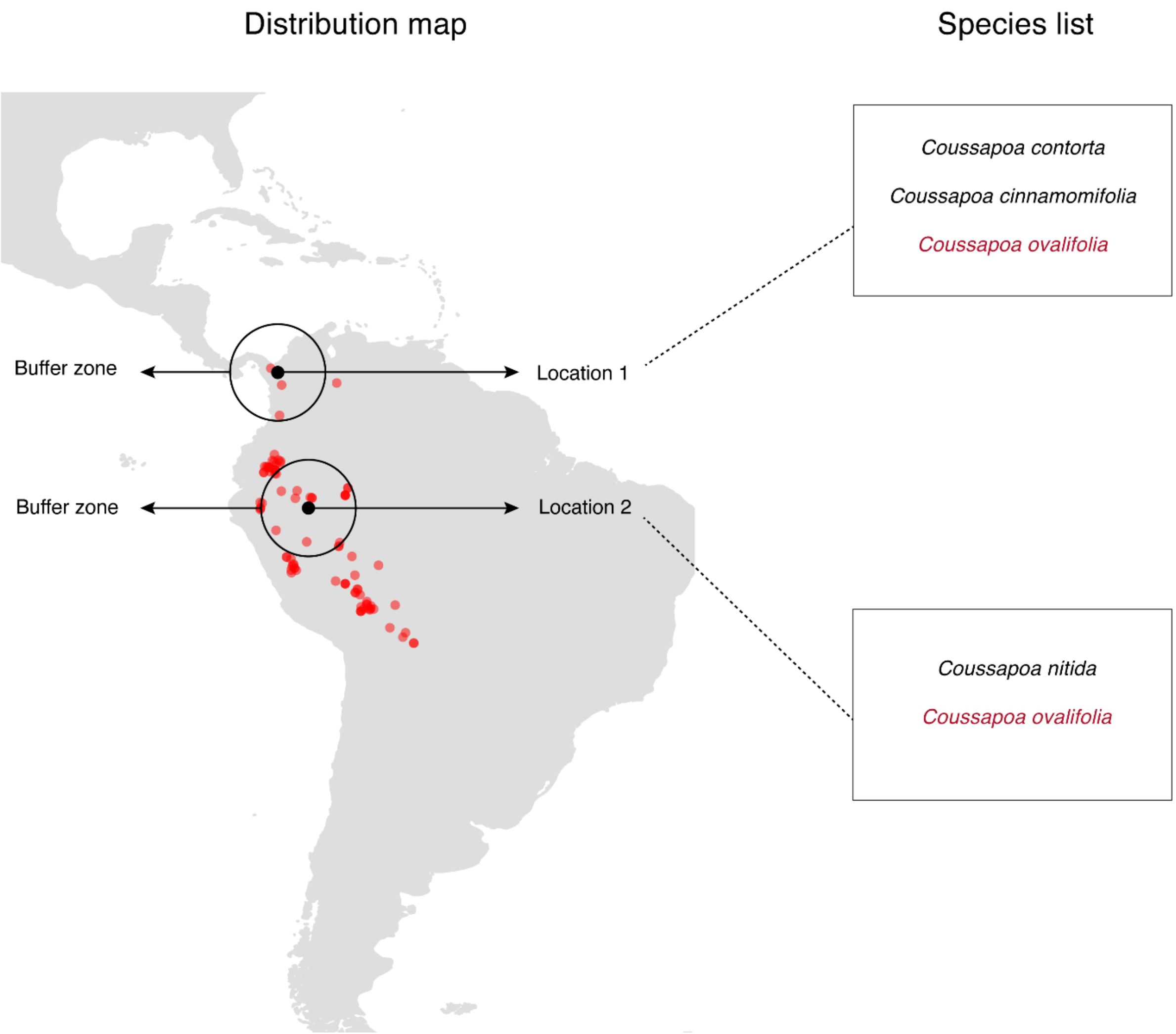


**Fig. S6. Graphical example of step 6 of the plant species cleaning process.** In this example, there are two occurrences of species labelled '*Coussapoa* sp.' in separate studies (locations 1 and 2). The distribution of *Coussapoa ovalifolia* (red dots) simultaneously overlaps the buffer zones of two 'problematic species' (i.e., species with genus name only) belonging to the same genus, such that these problematic species could not confidently be considered as being separate species. A distribution map like this was created for all congeneric species with occurrence data in either buffer zone. Note that *C. ovalifolia* is present in the potential list of *Coussapoa* species in both network locations (other *Coussapoa* species were omitted in the species lists for clarity).

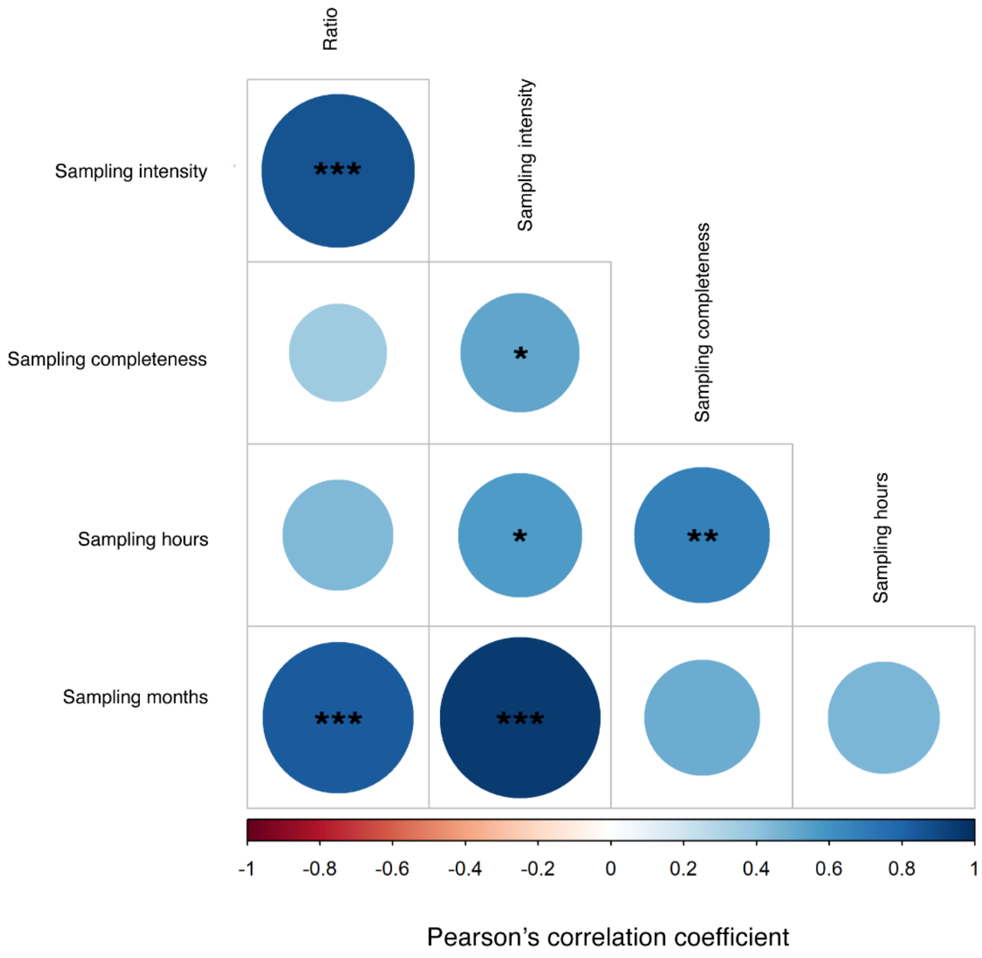


**Fig. S7. Results from Pearson’s correlation tests between sampling and network metrics.** For this analysis, we used the subset of networks sampled in Aotearoa New Zealand (n = 14). Asterisks represent significant correlations [*(P = 0.05 - 0.01), **(P = 0.01 - 0.001), ***(P < 0.001)]. Sizes of the circles are proportional to the correlation coefficient.

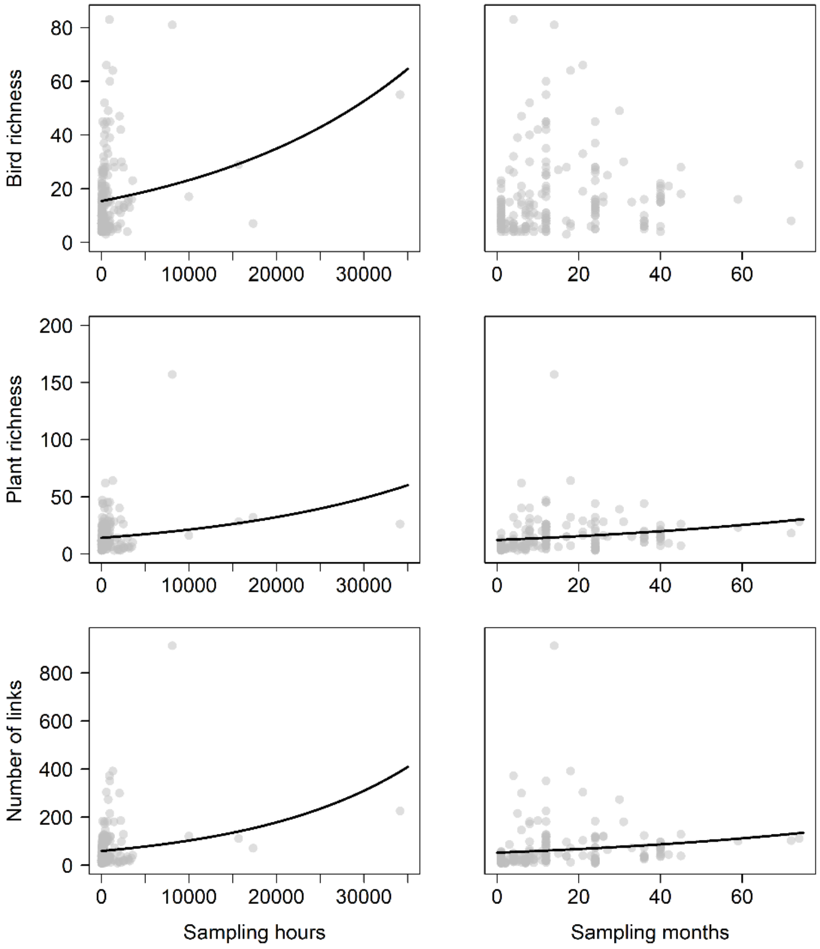


**Fig. S8. Relationships between network metrics and sampling hours and months.** We used generalized linear models (with Poisson errors, fitted with quasi-likelihood to deal with overdispersion) to obtain significance values. Points represent the 196 local frugivory networks in our dataset. Solid lines represent significant relationships.

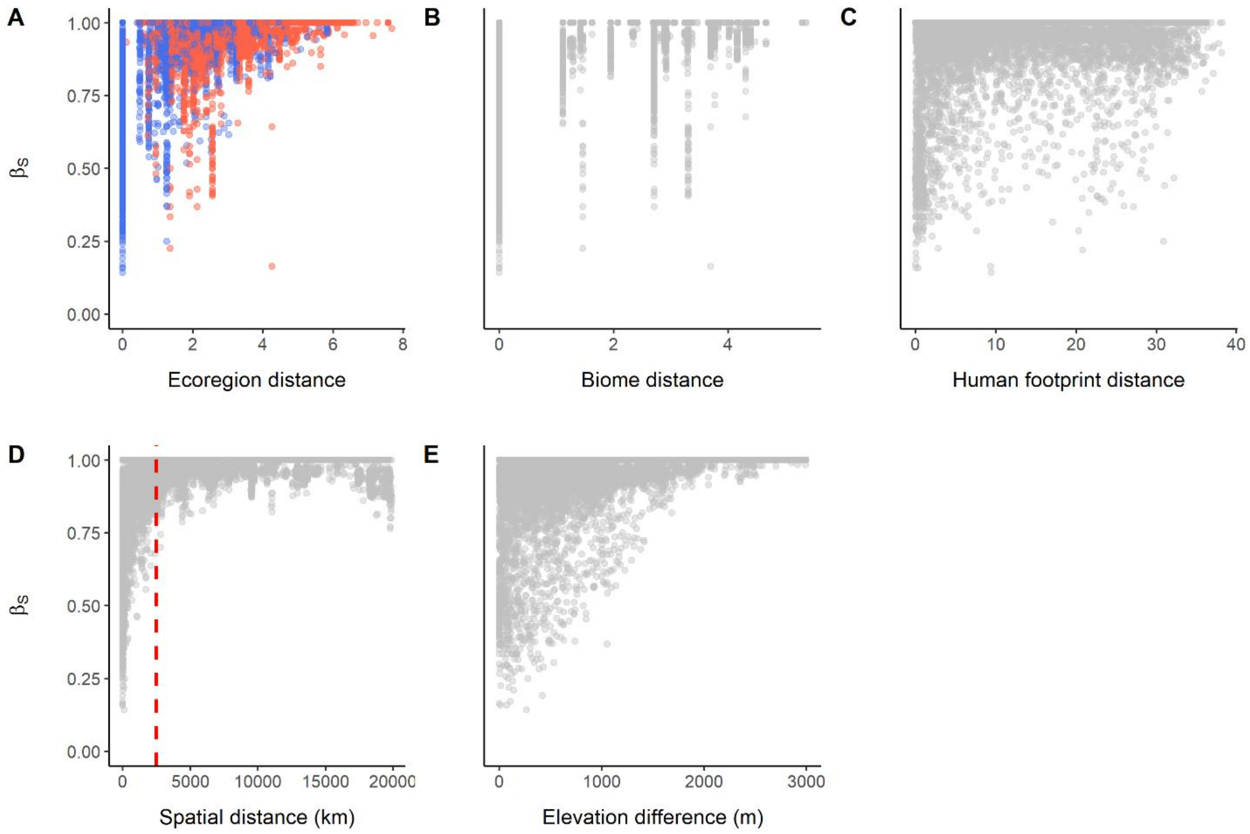


**Fig. S9. Scatterplots of the relationships between our predictor variables of interest (not those used for controlling sampling effects) and species turnover ($\beta_S$).** (**A**) The relationship between the quantitative version (environmental dissimilarity) of ecoregion distance and species turnover; point colors indicate whether the pair of local networks belong to the same (blue) or distinct (red) biomes. (**B**) The relationship between the quantitative version (environmental dissimilarity) of biome distance and species turnover. (**C**) The relationship between human footprint distance and species turnover. **(D)** The relationship between spatial distance and species turnover. Note that, contrary to species interactions (Figs. 1 and S14C), several networks still shared species beyond the threshold distance of 2,500 km (dotted red line). (**E**) The relationship between elevation difference and species turnover.

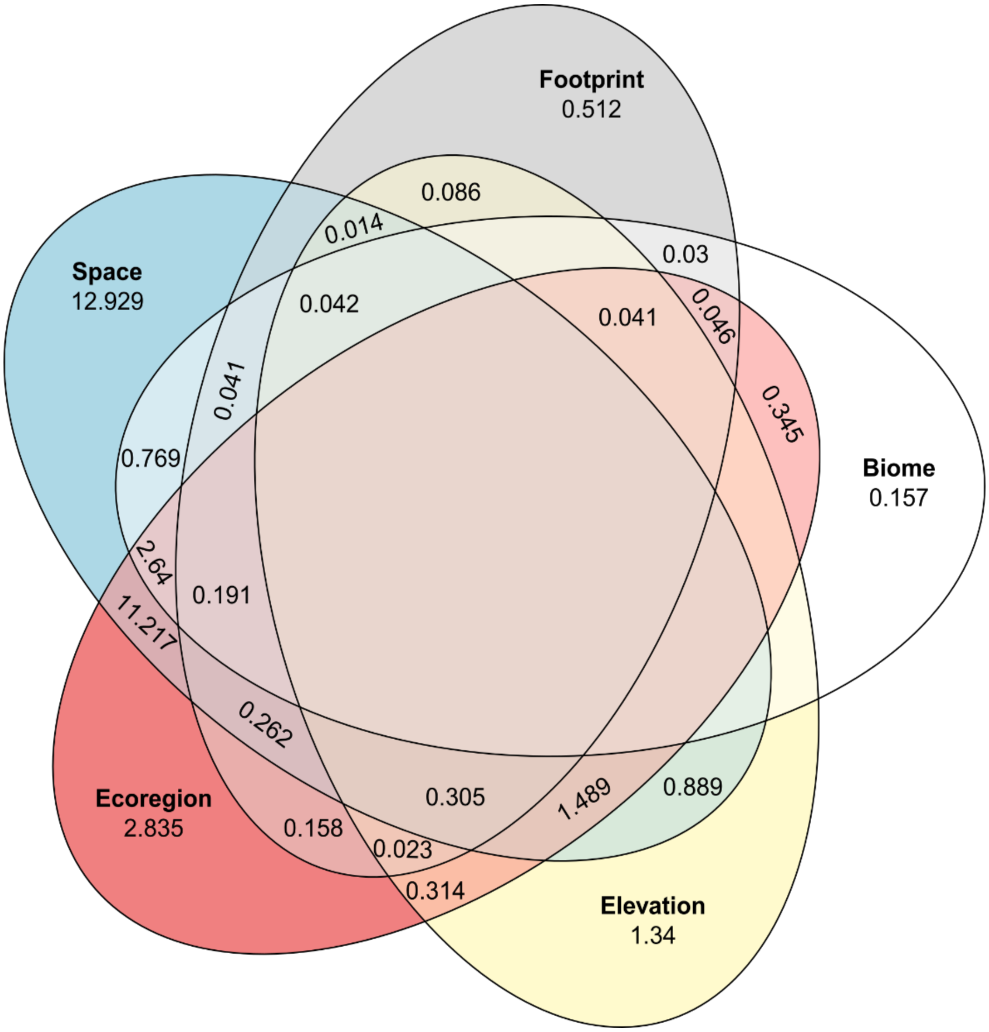


**Fig. S10. Venn diagram showing the relative contributions (%) of our main predictor variables to explaining the variation in species turnover ($\beta_S$) across networks, calculated using deviance partitioning.** Spatial distance alone explained the greatest proportion (12.9%) of the variation in species turnover, followed by the shared effect of spatial distance and ecoregion boundaries. Note that, to aid visualization, we only included our predictor variables of interest (i.e., not those used for controlling sampling effects). Terms that reduce explanatory power are not shown.

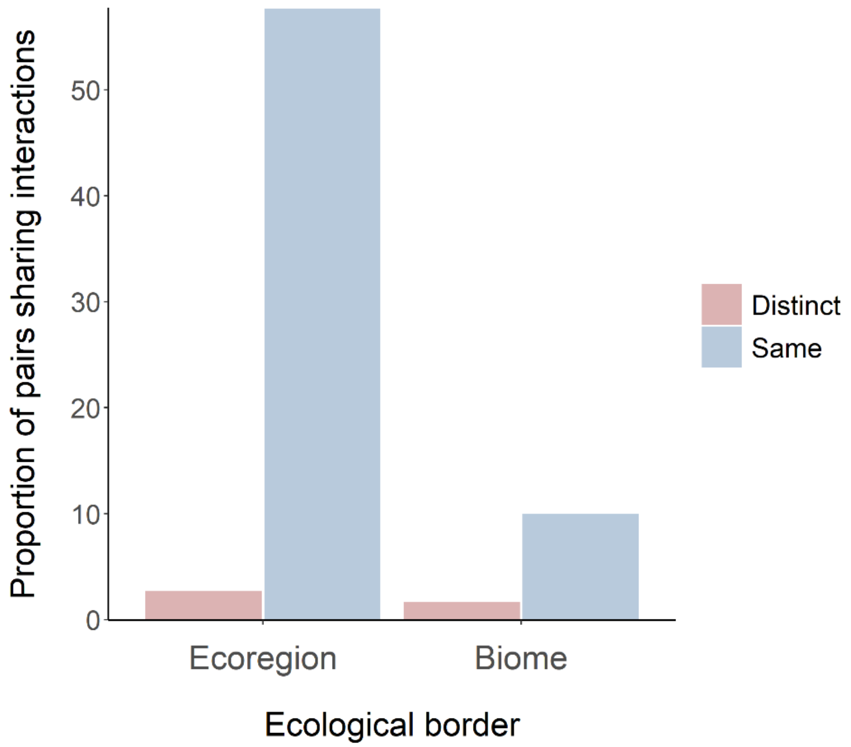


**Fig. S11. The effect of large-scale ecological boundaries on the proportion of pairs of local networks sharing interactions.** Avian frugivory networks located within the same ecoregion/biome were more likely to share interactions than those located across distinct ecoregions/biomes. Note that over half of the pairs of networks located within the same ecoregion shared at least one interaction.

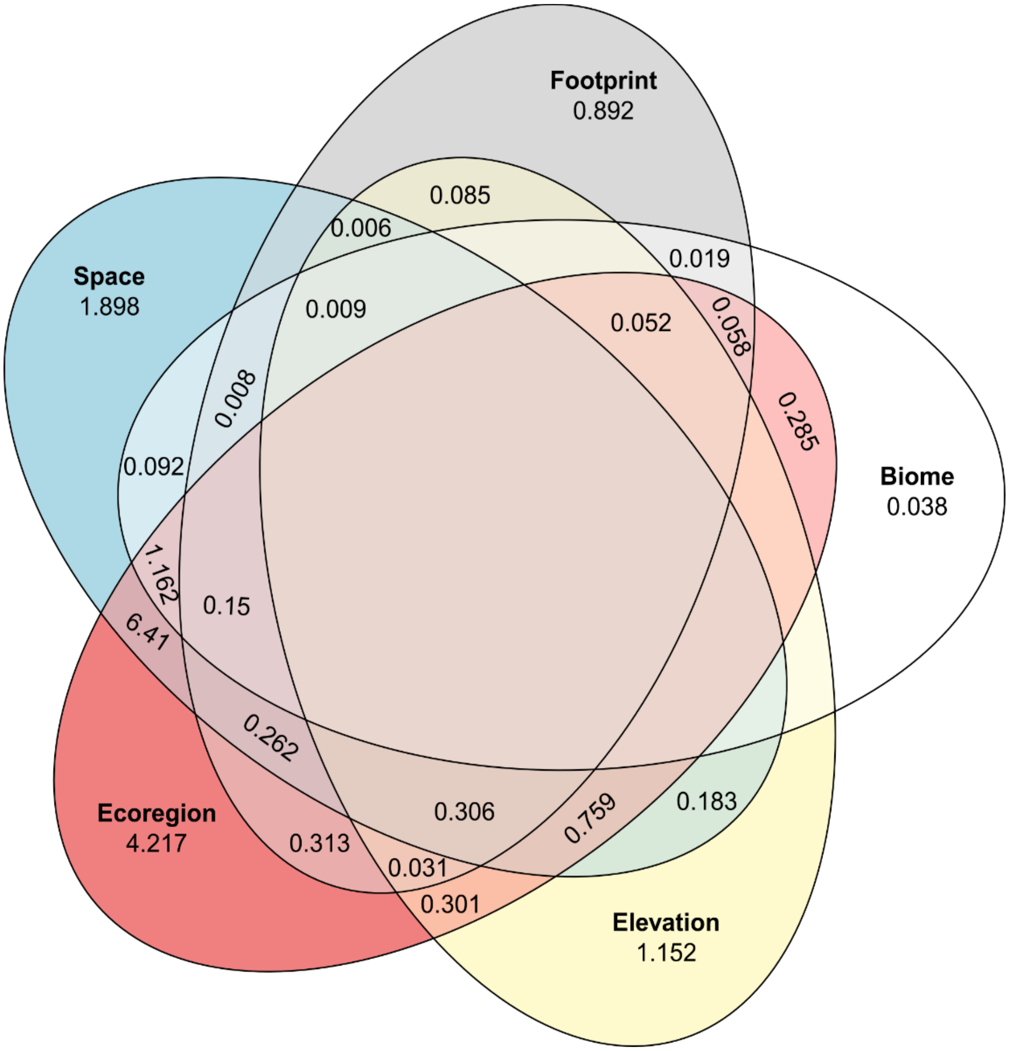


**Fig. S12. Venn diagram showing the relative contributions (%) of our main predictor variables to explaining the variation in plant-frugivore interaction dissimilarity ($\beta_{WN}$), calculated using deviance partitioning.** The shared effect of ecoregions and spatial distance explained the greatest proportion (6.41%) of the variation in interaction dissimilarity, followed by the unique contributions of these two variables. Note that, to aid visualization, we only included our predictor variables of interest (i.e., not those used for controlling sampling effects). Terms that reduce explanatory power are not shown.

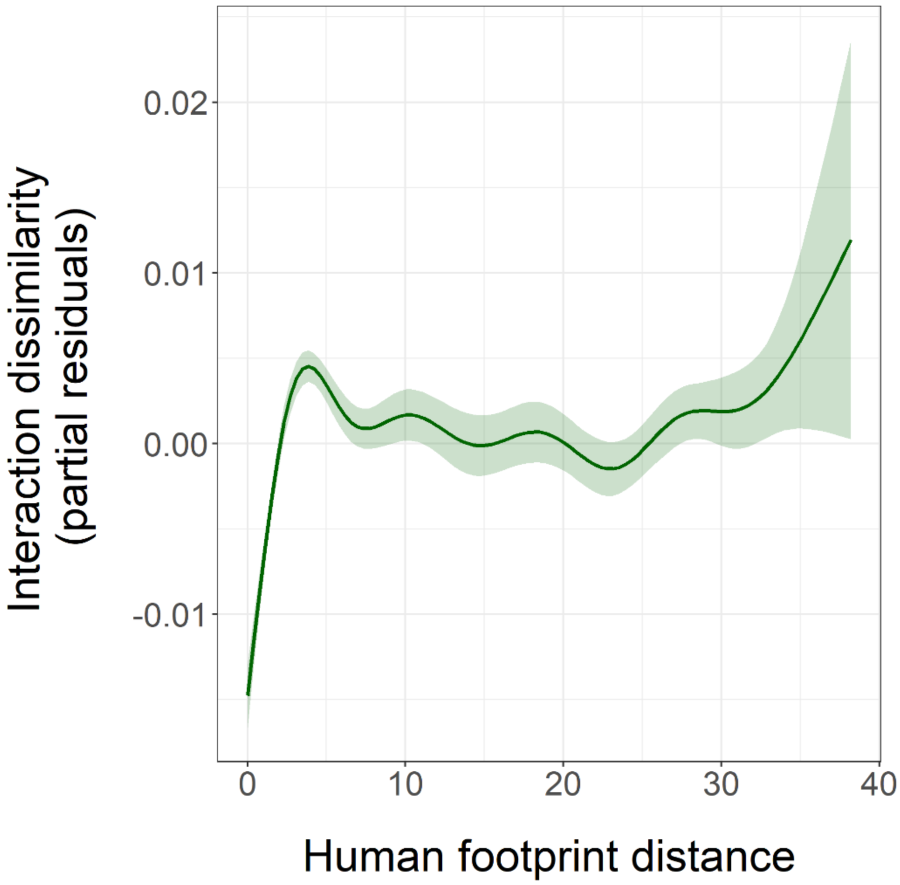


**Fig. S13. Partial effects plot of the relationship between human footprint distance and interaction dissimilarity ($\beta_{WN}$).** The smoothed line was fitted using a generalized additive model (GAM) with interaction dissimilarity as response variable and all of our predictor variables included (see Table 1). The lighter green area represents the 95% confidence interval of the fitted GAM.

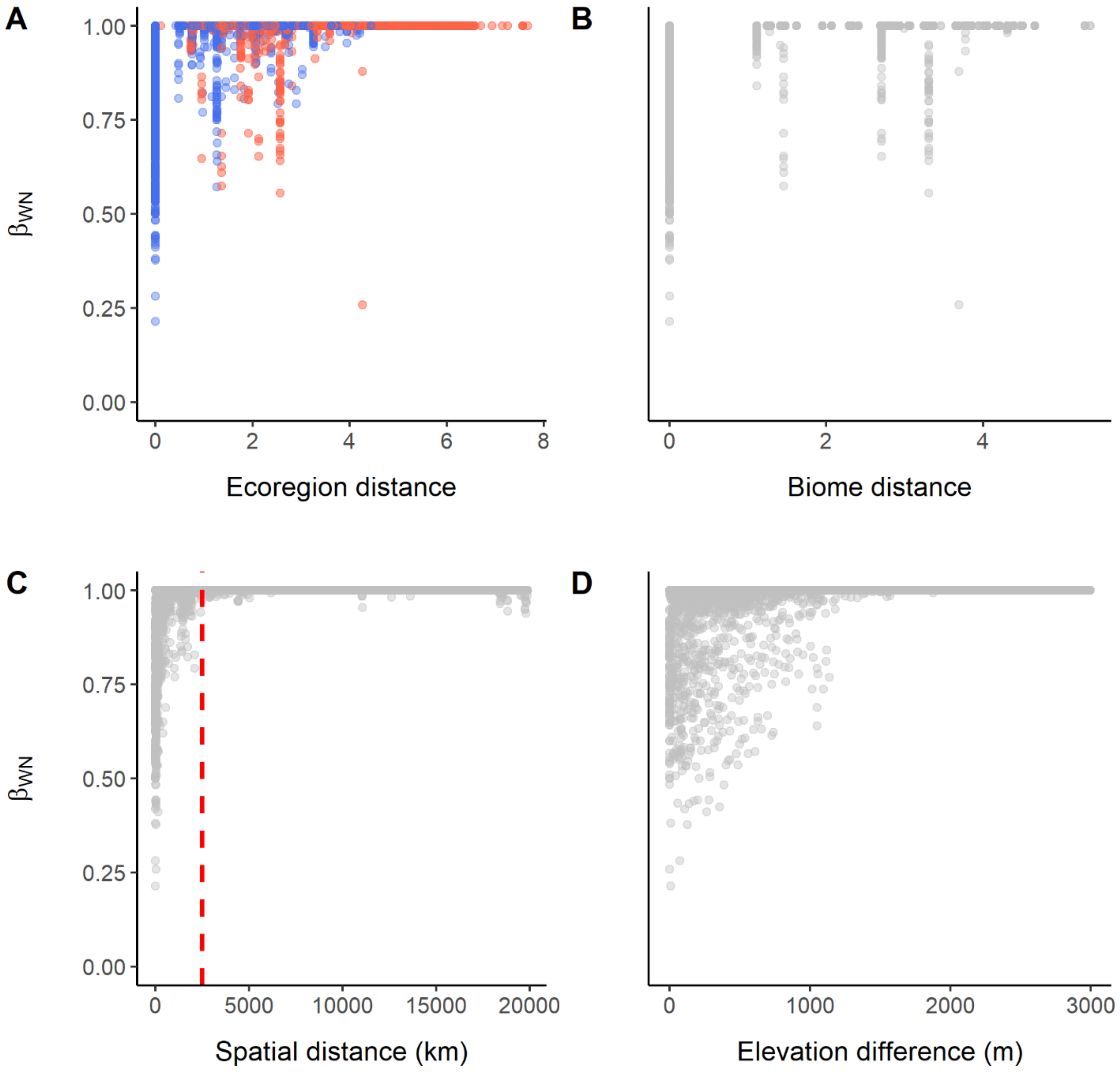


**Fig. S14. Scatterplots of the relationships between our predictor variables of interest (except human footprint distance, which is presented in the main text) and interaction dissimilarity ($\beta_{WN}$).** (**A**) The relationship between the quantitative version (environmental dissimilarity) of ecoregion distance and interaction dissimilarity; point colors indicate whether the pair of networks belong to the same (blue) or distinct (red) biomes. (**B**) The relationship between the quantitative version (environmental dissimilarity) of biome distance and interaction dissimilarity. (**C**) The relationship between spatial distance and interaction dissimilarity. Note that interaction dissimilarity increases sharply until a threshold distance of 2,500 km (dotted red line), beyond which few networks shared interactions (a similar pattern can be seen in Fig. 5). (**D**) The relationship between elevation difference and interaction dissimilarity.

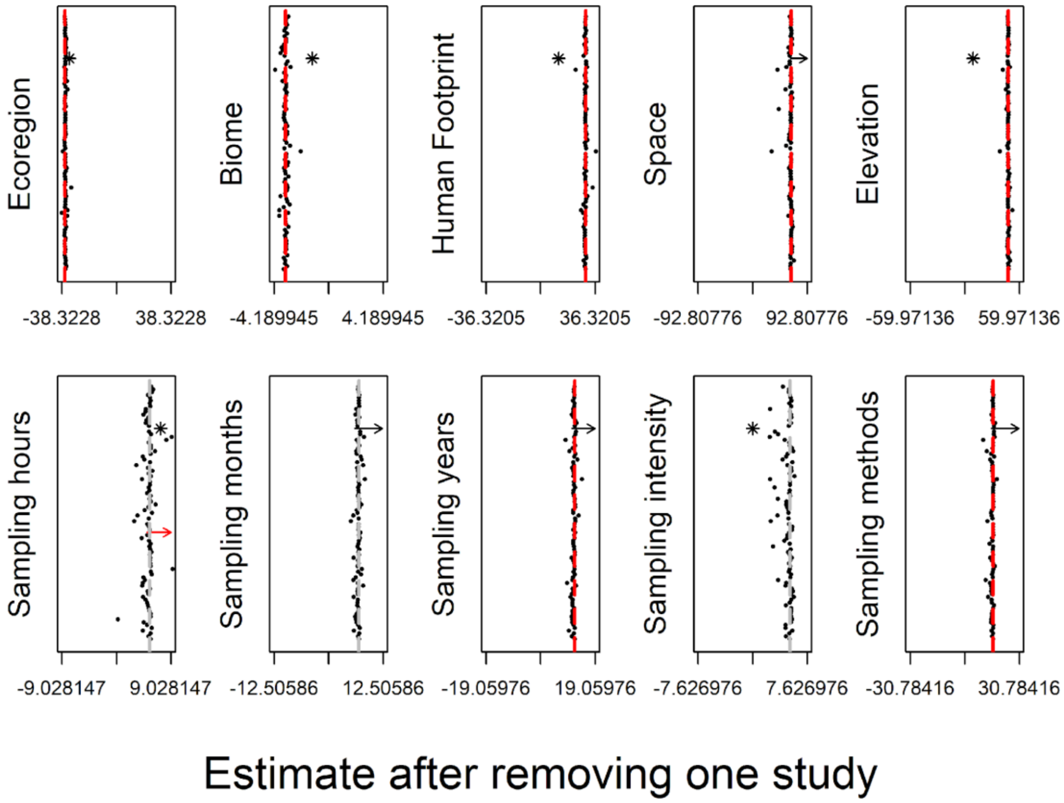


**Fig. S15. Effect of individual studies on estimates of t (for ecoregion and biome) and F values (for the remaining predictor variables) of generalized additive models with interaction dissimilarity ($\beta_{WN}$) as response variable.** Points represent estimate values after removing one study from the data, while asterisks indicate the estimates when the study with the greatest number of networks (n = 35) in our dataset, study 76 (*79*), is removed from the data. The estimates of the full model (with all studies included) are represented by the vertical lines. Red lines indicate a significant effect ($P < 0.05$), while gray lines indicate a non-significant effect. P values were calculated using a combination of generalized additive models and multiple regression on distance matrices (see ‘Statistical analyses’ section). The range of the x-axis was defined as ± 3 times the standard deviation of the estimates. Arrows indicate outliers beyond this range (black: when study 76 is removed; red: when other studies are removed).

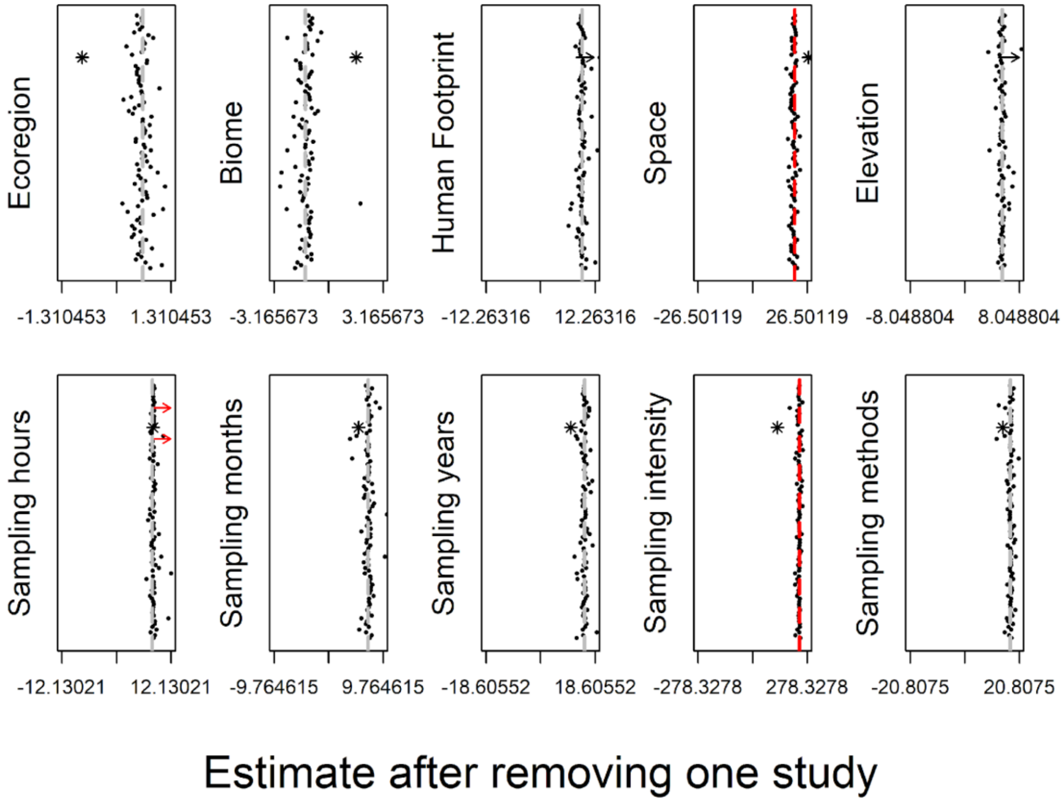


**Fig. S16. Effect of individual studies on estimates of t (for ecoregion and biome) and F values (for the remaining predictor variables) of generalized additive models with network structural dissimilarity as response variable.** Points represent estimate values after removing one study from the data, while asterisks indicate the estimates when the study with the greatest number of networks (n = 35) in our dataset, study 76 (*79*), is removed from the data. The estimates of the full model (with all studies included) are represented by the vertical lines. Red lines indicate a significant effect ($P < 0.05$), while gray lines indicate a non-significant effect. P values were calculated using a combination of generalized additive models and multiple regression on distance matrices (see ‘Statistical analyses’ section). The range of the x-axis was defined as ± 3 times the standard deviation of the estimates. Arrows indicate outliers beyond this range (black: when study 76 is removed; red: when other studies are removed).

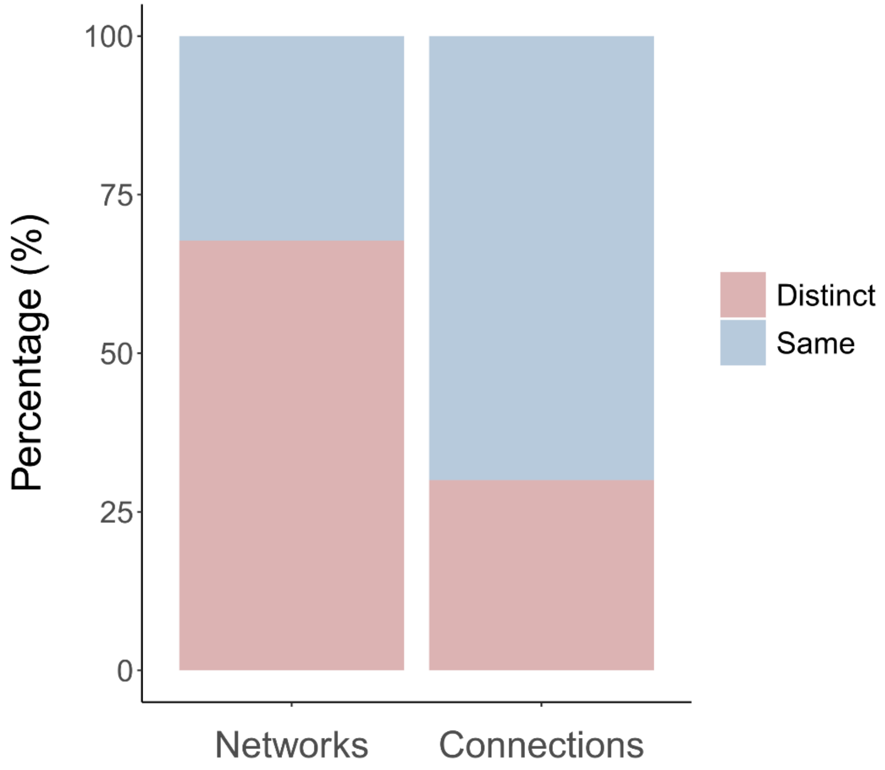


**Fig. S17. Percentage of long-distance network comparisons and connections (shared interactions) across ('distinct') and within ('same') biomes.** Around 67% of the long-distance network comparisons involved networks from distinct biomes, while most long-distance connections (70%) involved networks from the same biome.

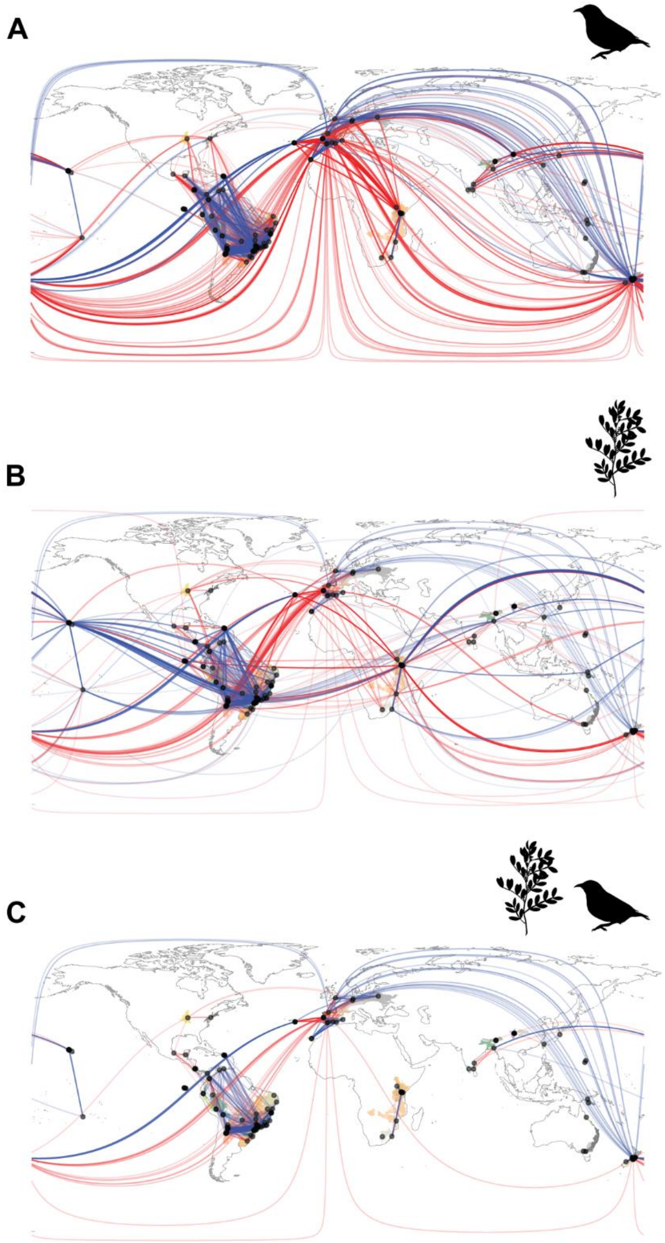


**Fig. S18. Plant and bird species connecting local networks, ecoregions and biomes.** World map with points representing the 196 local avian frugivory networks in our dataset. Colors of shaded areas represent the 67 ecoregions where networks were located, with similar colors indicating ecoregions that belong to the same biome. Lines represent the connections (shared species) plotted along the great circle distance between networks. Blue lines represent connections within biomes, while red lines represent connections across biomes. Stronger colour tones of lines indicate higher similarity of species ($1-\beta_S$) between networks. (**A**) Lines represent connections between networks sharing frugivorous bird species. (**B**) Lines represent connections between networks sharing plant species. (**C**) Lines represent connections between networks sharing both plant and frugivorous bird species (see Fig. 1 for the world map of shared plant-frugivore interactions).

**Table S1. Description of the 196 avian frugivory networks in our dataset.** Geographic coordinates were rounded to two decimal places. The metadata of each network (e.g., original coordinates, sampling methods, ecoregion, biome) is available at Dryad (*34*).

| Network ID | Latitude | Longitude | Location | Reference |
|---|---|---|---|---|
| 1* | 40.33 | -74.67 | New Jersey, USA | *82* |
| 2* | 18.30 | -66.78 | Caguana, Puerto Rico | *83* |
| 3* | 18.26 | -66.53 | Cialitos, Puerto Rico | *83* |
| 4* | 18.17 | -66.59 | Cordillera, Puerto Rico | *83* |
| 5* | 18.31 | -66.56 | Fronton, Puerto Rico | *83* |
| 6* | -28.95 | 31.75 | Mtunzini, South Africa | *84* |
| 7* | -22.82 | -47.11 | Mata Santa Genebra, São Paulo, Brazil | *85* |
| 8* | -22.82 | -47.11 | Mata Santa Genebra, São Paulo, Brazil | *85* |
| 9* | 18.51 | -89.49 | Campeche state, Mexico | *86* |
| 10* | 51.77 | -1.33 | Oxford, United Kingdom | *87* |
| 11* | -24.32 | -48.39 | Intervales, São Paulo, Brazil | *88* |
| 12 | -24.13 | -47.95 | Carlos Botelho, São Paulo, Brazil | *89* |
| 13 | -25.13 | -47.96 | Ilha do Cardoso, São Paulo, Brazil | *90* |
| 14 | -22.55 | -42.28 | Poço das Antas, Rio de Janeiro, Brazil | *91* |
| 15 | -23.55 | -45.06 | Ilha Anchieta, São Paulo, Brazil | *92* |
| 16 | -20.80 | -42.86 | Viçosa, Minas Gerais, Brazil | *93* |
| 17 | -28.22 | -51.17 | Estação Aracuri, Rio Grande do Sul, Brazil | *94* |
| 18 | -22.94 | -46.75 | Itatiba, São Paulo, Brazil | *95* |
| 19 | -22.48 | -47.59 | Rio Claro, São Paulo, Brazil | *96* |
| 20 | -22.82 | -47.43 | Santa Barbara do Oeste, São Paulo, Brazil | *97* |
| 21 | -22.67 | -47.20 | Cosmópolis, São Paulo, Brazil | *97* |
| 22 | -22.57 | -47.50 | Iracemápolis, São Paulo, Brazil | *97* |
| 23 | -23.55 | -46.72 | São Paulo, Brazil | *98* |
| 24 | -22.71 | -47.61 | Piracicaba, São Paulo, Brazil | *99* |
| 25 | -22.77 | -43.69 | Rio de Janeiro, Brazil | *100* |
| 26 | 37.79 | -25.18 | Azores, Portugal | *101* |
| 27* | 0.30 | 34.79 | Kakamega Forest, Kenya | *102* |
| 28 | -25.49 | -49.26 | Curitiba, Paraná, Brazil | *103* |
| 29 | -25.44 | -49.24 | Curitiba, Paraná, Brazil | *103* |
| 30 | -25.44 | -49.22 | Curitiba, Paraná, Brazil | *103* |
| 31 | -25.42 | -49.37 | Curitiba, Paraná, Brazil | *103* |
| 32 | -25.41 | -49.27 | Curitiba, Paraná, Brazil | *103* |
| 33 | -25.36 | -49.26 | Curitiba, Paraná, Brazil | *103* |
| 34 | -25.38 | -49.32 | Curitiba, Paraná, Brazil | *103* |
| 35 | -25.17 | -48.41 | Paraná, Brazil | *104* |
| 36 | 28.03 | -15.46 | Bandama, Gran Canaria, Spain | *105* |
| 37 | 28.07 | -15.46 | El Palomar, Gran Canaria, Spain | *105* |
| 38 | -12.99 | -41.34 | Chapada Diamantina, Bahia, Brazil | *106* |
| 39 | 37.18 | -6.32 | Hato Ratón, Sevilla, Spain | *107* |
| 40 | -16.40 | -67.50 | Chulumani, Bolivia | *108* |
| 41 | 30.33 | 130.50 | Yakushima Island, Japan | *109* |

| Network ID | Latitude | Longitude | Location | Reference |
|---|---|---|---|---|
| 42 | -18.95 | -48.20 | Uberlândia, Minas Gerais, Brazil | *110* |
| 43 | 21.44 | -158.08 | Ēkahanui, Hawai'i, USA | *28* |
| 44 | 21.54 | -158.19 | Kahanahāiki, Hawai'i, USA | *28* |
| 45 | 21.38 | -157.87 | Moanalua, Hawai'i, USA | *28* |
| 46 | 21.51 | -158.14 | Mount Ka'ala, Hawai'i, USA | *28* |
| 47 | 21.54 | -158.18 | Pahole, Hawai'i, USA | *28* |
| 48 | 21.34 | -157.81 | Tantalus, Hawai'i, USA | *28* |
| 49 | 21.63 | -158.04 | Waimea Valley, Hawai'i, USA | *28* |
| 50 | 37.57 | -0.91 | Sierra de la Fausilla, Murcia, Spain | *111* |
| 51 | 26.99 | 92.94 | Pakke Tiger Reserve, India | *112* |
| 52 | 7.77 | -76.67 | Tulenapa, Antioquia, Colombia | *113* |
| 53 | 43.28 | -5.50 | Cantabrian Range, Spain | *114* |
| 54 | -29.06 | -50.07 | Rio Grande do Sul, Brazil | *115* |
| 55 | -31.67 | -53.25 | Rio Grande do Sul, Brazil | *115* |
| 56 | 15.17 | 145.77 | Saipan, Mariana Islands | *116* |
| 57 | 14.14 | 145.21 | Rota, Mariana Islands | *116* |
| 58 | -0.75 | -90.32 | Santa Cruz, Galapagos Islands | *117* |
| 59 | 52.74 | 23.78 | Białowieża Forest, Poland | *118* |
| 60 | -4.92 | -73.75 | Jenaro Herrera, Peru | *119* |
| 61 | 18.47 | -67.11 | Finca Montaña, Aguadilla, Puerto Rico | *120* |
| 62 | 19.59 | -96.38 | Veracruz, Mexico | *121* |
| 63 | -8.97 | -36.05 | Coimbra Forest, Alagoas, Brazil | *122* |
| 64 | -41.29 | 174.73 | Wellington, Aotearoa New Zealand | *30* |
| 65 | -41.29 | 174.75 | Wellington, Aotearoa New Zealand | *30* |
| 66 | -41.30 | 174.75 | George Denton Park, Aotearoa New Zealand | *30* |
| 67 | -41.29 | 174.80 | Charles Plimmer Park, Aotearoa New Zealand | *30* |
| 68 | -41.28 | 174.77 | Wellington, Aotearoa New Zealand | *30* |
| 69 | -42.35 | 173.57 | Hinau Reserve, Aotearoa New Zealand | *30* |
| 70 | -42.33 | 173.63 | Mount Fyffe Reserve, Aotearoa New Zealand | *30* |
| 71 | -42.28 | 173.74 | Puhi-Puhi, Aotearoa New Zealand | *30* |
| 72 | -42.24 | 173.78 | Blue Duck Reserve, Aotearoa New Zealand | *30* |
| 73 | 40.22 | -8.46 | Choupal, Coimbra, Portugal | *123* |
| 74 | -41.30 | 174.75 | Wellington, Aotearoa New Zealand | *124* |
| 75 | -12.93 | -38.40 | Salvador, Bahia, Brazil | *125* |
| 76 | 26.93 | 92.97 | Pakke Tiger Reserve, India | *126* |
| 77 | 27.02 | 92.95 | Papum Reserve Forest, India | *126* |
| 78 | -43.75 | 169.40 | Windbag Valley, Aotearoa New Zealand | *127* |
| 79 | 37.78 | -25.15 | Azores, Portugal | *101* |
| 80 | 37.80 | -25.16 | Azores, Portugal | *101* |
| 81 | 37.79 | -25.16 | Azores, Portugal | *101* |
| 82 | 40.31 | -8.40 | Coimbra, Portugal | *128* |
| 83 | 40.26 | -8.48 | Coimbra, Portugal | Unpublished |
| 84 | -0.66 | -90.32 | Santa Cruz, Galapagos Islands | *117* |
| 85 | -0.91 | -89.43 | San Cristóbal, Galapagos Islands | *117* |
| 86 | -0.89 | -89.49 | San Cristóbal, Galapagos Islands | *117* |

| Network ID | Latitude | Longitude | Location | Reference |
|---|---|---|---|---|
| 87 | -19.95 | 34.37 | Gorongosa National Park, Mozambique | *129* |
| 88 | 13.70 | 80.19 | Sriharikota Island, India | *130* |
| 89 | -5.05 | -37.52 | Furna Feia, Rio Grande do Norte, Brazil | *131* |
| 90 | -7.22 | 146.81 | Mount Missim, New Guinea | *132* |
| 91 | -9.45 | 147.35 | Varirata National Park, New Guinea | *133* |
| 92 | -29.12 | 26.17 | Bloemfontein, South Africa | *134* |
| 93 | -37.62 | 144.42 | Lerderderg Park, Australia | *135* |
| 94 | -37.72 | 145.57 | Mt Healesville and Donna Buang, Australia | *136* |
| 95 | -41.33 | 173.05 | Brightwater, Aotearoa New Zealand | *137* |
| 96 | -41.32 | 173.26 | Nelson, Aotearoa New Zealand | *137* |
| 97 | -41.41 | 173.04 | Faulkners, Wakefield, Aotearoa New Zealand | *137* |
| 98 | -22.28 | -41.68 | Restinga de Jurubatiba, Rio de Janeiro, Brazil | *138* |
| 99 | -15.95 | -47.97 | Brasília, Brazil | *139* |
| 100 | -19.77 | -40.04 | Comboios, Espírito Santo, Brazil | *140* |
| 101 | -23.37 | -46.60 | Cantareira, São Paulo, Brazil | *141* |
| 102 | -19.57 | -56.20 | Pantanal, Brazil | *142* |
| 103 | -22.39 | -47.54 | Rio Claro, São Paulo, Brazil | Unpublished |
| 104 | -21.73 | -48.02 | Araraquara, São Paulo, Brazil | *143* |
| 105 | -24.73 | -64.67 | El Rey National Park, Argentina | *144* |
| 106 | -27.25 | -65.88 | Campo de Los Alisos, Argentina | *144* |
| 107 | -27.23 | -65.62 | La Florida Provincial Park, Argentina | *144* |
| 108 | -26.80 | -65.30 | San Javier y Yerba Huasi, Argentina | *145* |
| 109 | -24.76 | -64.69 | Pozo Verde, El Rey National Park, Argentina | *146* |
| 110 | -27.03 | -65.77 | Quebrada del Portugues, Argentina | *144* |
| 111 | -24.10 | -64.45 | EcoPortal de Piedra, Argentina | *144* |
| 112 | -23.69 | -64.88 | Calilegua National Park, Argentina | *144* |
| 113 | -23.69 | -64.87 | Calilegua National Park, Argentina | *144* |
| 114 | -22.28 | -64.71 | El Nogalar de los Toldos, Argentina | *144* |
| 115 | -26.75 | -65.33 | Parque Sierra de San Javier, Argentina | *146* |
| 116 | -26.80 | -65.33 | Parque Sierra de San Javier, Argentina | Unpublished |
| 117 | -15.35 | -39.20 | Bahia, Brazil | *147* |
| 118 | -15.21 | -39.14 | Bahia, Brazil | *147* |
| 119 | -15.13 | -39.12 | Bahia, Brazil | *147* |
| 120 | -15.25 | -39.08 | Bahia, Brazil | *147* |
| 121 | -15.26 | -39.09 | Bahia, Brazil | *147* |
| 122 | 10.28 | -84.05 | Rara Avis Reserve, Costa Rica | *148* |
| 123 | -17.85 | 146.08 | Mission Beach, Queensland, Australia | *149* |
| 124 | 10.35 | 77.04 | Valparai and Anamalai Reserve, India | *150* |
| 125 | 31.07 | 103.71 | Dujiangyan, Sichuan Province, China | *151* |
| 126 | 31.05 | 103.74 | Dujiangyan, Sichuan Province, China | *151* |
| 127 | 31.05 | 103.73 | Dujiangyan, Sichuan Province, China | *151* |
| 128 | 31.06 | 103.72 | Dujiangyan, Sichuan Province, China | *151* |
| 129 | 31.05 | 103.72 | Dujiangyan, Sichuan Province, China | *151* |
| 130 | 31.08 | 103.70 | Dujiangyan, Sichuan Province, China | *151* |
| 131 | 31.09 | 103.72 | Dujiangyan, Sichuan Province, China | *151* |

| Network ID | Latitude | Longitude | Location | Reference |
|---|---|---|---|---|
| 132 | 31.09 | 103.73 | Dujiangyan, Sichuan Province, China | *151* |
| 133 | 31.08 | 103.72 | Dujiangyan, Sichuan Province, China | *151* |
| 134 | 31.06 | 103.73 | Dujiangyan, Sichuan Province, China | *151* |
| 135 | 31.06 | 103.72 | Dujiangyan, Sichuan Province, China | *151* |
| 136 | 31.05 | 103.73 | Dujiangyan, Sichuan Province, China | *151* |
| 137 | 31.05 | 103.73 | Dujiangyan, Sichuan Province, China | *151* |
| 138 | 37.98 | -2.90 | Serranía de Cazorla, Spain | *152* |
| 139 | 37.38 | -5.71 | El Viso del Alcor, Sevilla, Spain | *152* |
| 140 | 50.30 | 8.66 | Friedberg, Hesse, Germany | *153* |
| 141 | 51.15 | 9.00 | Kellerwald-Edersee, Germany | *154* |
| 142 | -3.23 | 37.27 | Mt Kilimanjaro, Tanzania | *79* |
| 143 | -3.25 | 37.32 | Mt Kilimanjaro, Tanzania | *79* |
| 144 | -3.27 | 37.47 | Mt Kilimanjaro, Tanzania | *79* |
| 145 | -3.17 | 37.24 | Mt Kilimanjaro, Tanzania | *79* |
| 146 | -3.21 | 37.34 | Mt Kilimanjaro, Tanzania | *79* |
| 147 | -3.26 | 37.42 | Mt Kilimanjaro, Tanzania | *79* |
| 148 | -3.26 | 37.42 | Mt Kilimanjaro, Tanzania | *79* |
| 149 | -3.23 | 37.52 | Mt Kilimanjaro, Tanzania | *79* |
| 150 | -3.14 | 37.24 | Mt Kilimanjaro, Tanzania | *79* |
| 151 | -3.13 | 37.24 | Mt Kilimanjaro, Tanzania | *79* |
| 152 | -3.14 | 37.30 | Mt Kilimanjaro, Tanzania | *79* |
| 153 | -3.14 | 37.31 | Mt Kilimanjaro, Tanzania | *79* |
| 154 | -3.17 | 37.36 | Mt Kilimanjaro, Tanzania | *79* |
| 155 | -3.15 | 37.29 | Mt Kilimanjaro, Tanzania | *79* |
| 156 | -3.18 | 37.36 | Mt Kilimanjaro, Tanzania | *79* |
| 157 | -3.19 | 37.51 | Mt Kilimanjaro, Tanzania | *79* |
| 158 | -3.20 | 37.52 | Mt Kilimanjaro, Tanzania | *79* |
| 159 | -3.19 | 37.44 | Mt Kilimanjaro, Tanzania | *79* |
| 160 | -3.10 | 37.26 | Mt Kilimanjaro, Tanzania | *79* |
| 161 | -3.17 | 37.36 | Mt Kilimanjaro, Tanzania | *79* |
| 162 | -3.16 | 37.36 | Mt Kilimanjaro, Tanzania | *79* |
| 163 | -3.19 | 37.44 | Mt Kilimanjaro, Tanzania | *79* |
| 164 | -3.18 | 37.51 | Mt Kilimanjaro, Tanzania | *79* |
| 165 | -3.18 | 37.25 | Mt Kilimanjaro, Tanzania | *79* |
| 166 | -3.30 | 37.50 | Mt Kilimanjaro, Tanzania | *79* |
| 167 | -3.33 | 37.50 | Mt Kilimanjaro, Tanzania | *79* |
| 168 | -3.30 | 37.62 | Mt Kilimanjaro, Tanzania | *79* |
| 169 | -3.19 | 37.25 | Mt Kilimanjaro, Tanzania | *79* |
| 170 | -3.27 | 37.60 | Mt Kilimanjaro, Tanzania | *79* |
| 171 | -3.32 | 37.67 | Mt Kilimanjaro, Tanzania | *79* |
| 172 | -3.37 | 37.45 | Mt Kilimanjaro, Tanzania | *79* |
| 173 | -3.38 | 37.50 | Mt Kilimanjaro, Tanzania | *79* |
| 174 | -3.33 | 37.64 | Mt Kilimanjaro, Tanzania | *82* |
| 175 | -3.32 | 37.68 | Mt Kilimanjaro, Tanzania | *82* |
| 176 | -3.31 | 37.68 | Mt Kilimanjaro, Tanzania | *79* |

| Network ID | Latitude | Longitude | Location | Reference |
|---|---|---|---|---|
| 177 | -16.40 | -67.50 | Chulumani, Bolivia | *108* |
| 178 | 4.72 | -75.57 | Otún Quimbaya, Colombia | *155* |
| 179 | 4.70 | -75.48 | Ucumarí, Colombia | *155* |
| 180 | -3.96 | -79.06 | Podocarpus National Park, Ecuador | *19* |
| 181 | -4.10 | -79.17 | Podocarpus National Park, Ecuador | *19* |
| 182 | -13.05 | -71.54 | San Pedro, Peru | *156* |
| 183 | -13.17 | -71.58 | Wayqecha, Peru | *156* |
| 184 | 9.71 | -69.58 | Yacambú National Park, Venezuela | *157* |
| 185 | 10.39 | -67.02 | Altos de Pipe, Coastal Cordillera, Venezuela | *157* |
| 186 | 10.30 | 79.85 | Point Calimere Wildlife Sanctuary, India | *158* |
| 187 | 20.60 | -156.33 | Kanaio Natural Area Reserve, Hawai'i | *159* |
| 188 | -3.37 | 38.33 | Taita Hills, Kenya | *160* |
| 189 | 40.13 | -88.17 | Champaign County, Illinois, USA | 161 |
| 190 | -17.53 | -149.83 | Moorea, French Polynesia | *162* |
| 191 | 10.47 | -83.51 | Tortuguero, Costa Rica | *163* |
| 192 | 24.80 | 121.25 | Fushan Experimental Forest, Taiwan | *164* |
| 193 | 22.46 | 91.77 | Chittagong, Bangladesh | *165* |
| 194 | 10.42 | -84.01 | La Selva Biological Station, Costa Rica | *166* |
| 195 | 10.42 | -84.02 | La Selva Biological Station, Costa Rica | *167* |
| 196 | 39.14 | 2.94 | Cabrera Island, Spain | *168* |

* Obtained through the Web of Life database (*35*).

**Table S2. Quantitative metrics of network sampling.** Sampling intensity and completeness aim to account for how complete network sampling was in terms of species interactions, while sampling hours and months account for the time-span of the study (see more details on the 'Generating the distance matrices' section).

| Sampling metric | Rationale |
|---|---|
| Sampling intensity | Sampling intensity was calculated as the square-root of the number of interaction events divided by the square-root of the product of the number of plant and animal species in the local network [following (*169*)]. Sampling intensity was included in our models because it presented a strong and positive relationship with the ratio between the number of interactions sampled in the local network and the number of known possible interactions (among that same set of species) in the region (for the subset of networks within the Aotearoa New Zealand meta-network) (Fig. S7). |
| Sampling completeness | Sampling completeness was calculated as the observed richness of links divided by the estimated richness of links in the local network [following (*170*)]. We used the Chao 1 richness estimator (*171*) to obtain the estimated number of links in our networks. Sampling completeness was not included in our models because it did not present a significant relationship with the ratio between the number of interactions in the local network and the number of known possible interactions (among that same set of species) in the region (Fig. S7). Thus, we considered that this metric did not provide a good representation of how complete network sampling was in terms of species interactions. |
| Sampling hours | Number of sampling hours was included in our statistical models because it presented strong and positive relationships with bird richness, plant richness and number of links in the local networks (Fig. S8). |
| Sampling months | Number of sampling months was included in our statistical models because it presented a strong and positive relationship with the ratio between the number of interactions in the local network and the number of known possible interactions (among that same set of species) in the region (Fig. S7), as well as with plant richness and number of links in the local networks (for the entire dataset) (Fig. S8). |

**Table S3. Description of variables used to generate the methods dissimilarity matrix.**

| Variable | Description |
|---|---|
| Sampling design | Whether the sampling design was ‘transect’, ‘plot’, ‘mist-net’, ‘focal observation’, ‘camera-trap’, or any combination of these. |
| Sampling focus | Whether the focal organisms were frugivorous birds, plants, or both. |
| Sampling coverage | Whether there were focal species (‘partial coverage’) or not (‘total coverage’). |
| Interaction frequency type | Whether interaction frequency was estimated by counting the number of bird visits, number of fruits consumed by the bird, number of seeds in bird droppings, or number of bird droppings with seeds. |

**Table S4. Multiple drivers of species turnover (βs) on plant-frugivore networks.** Here, we used the binary version of ecoregion and biome distance matrices. *P* values were calculated using a combination of generalized additive models and multiple regression on distance matrices. EDF represents the estimated degrees of freedom for each smooth term in the model. Bold values indicate statistically significant results (*P* < 0.05). *N* pairs of networks = 19,110.

| **Parametric coefficients** | **Estimate** | **t** | **p** |
|---|---|---|---|
| Intercept | 0.976 | 1734.300 | **0.001** |
| Ecoregion (same) | -0.122 | -38.093 | **0.001** |
| Biome (same) | -0.008 | -8.799 | **0.001** |
| **Smooth Terms** | **EDF** | **F** | **p** |
| s (local footprint distance) | 8.312 | 28.504 | **0.001** |
| s (spatial distance) | 8.866 | 725.571 | **0.001** |
| s (elevational difference) | 5.589 | 99.954 | **0.001** |
| s (hours distance) | 6.917 | 4.004 | 0.619 |
| s (months distance) | 6.755 | 6.525 | 0.089 |
| s (years distance) | 6.402 | 7.422 | 0.068 |
| s (sampling intensity distance) | 1.007 | 26.580 | **0.005** |
| s (methods distance) | 8.039 | 10.911 | **0.015** |

**Table S5. Multiple drivers of species turnover (βs) on plant-frugivore networks.** Here, we used the quantitative version (environmental dissimilarity) of ecoregion and biome distance matrices. *P* values were calculated using a combination of generalized additive models and multiple regression on distance matrices. EDF represents the estimated degrees of freedom for each smooth term in the model. Bold values indicate statistically significant results ($P < 0.05$). *N* pairs of networks = 19,110.

| Smooth Terms | EDF | F | p |
|---|---|---|---|
| s (ecoregion distance) | 8.570 | 137.969 | **0.001** |
| s (biome distance) | 8.202 | 37.937 | **0.001** |
| s (local footprint distance) | 8.339 | 29.465 | **0.001** |
| s (spatial distance) | 8.890 | 698.382 | **0.001** |
| s (elevational difference) | 5.517 | 98.173 | **0.001** |
| s (hours distance) | 7.330 | 4.876 | 0.448 |
| s (months distance) | 5.371 | 5.811 | 0.109 |
| s (years distance) | 6.152 | 7.741 | 0.063 |
| s (sampling intensity distance) | 4.365 | 6.108 | 0.315 |
| s (methods distance) | 7.996 | 11.474 | **0.017** |

**Table S6. Multiple drivers of plant-frugivore interaction dissimilarity ($\beta_{WN}$).** Here, we used the quantitative version (environmental dissimilarity) of ecoregion and biome distance matrices. *P* values were calculated using a combination of generalized additive models and multiple regression on distance matrices. EDF represents the estimated degrees of freedom for each smooth term in the model. Bold values indicate statistically significant results ($P < 0.05$). *N* pairs of networks = 19,110.

| Smooth Terms | EDF | F | p |
|---|---|---|---|
| s (ecoregion distance) | 8.595 | 110.122 | **0.001** |
| s (biome distance) | 7.827 | 10.492 | **0.022** |
| s (local footprint distance) | 8.570 | 32.573 | **0.001** |
| s (spatial distance) | 8.855 | 81.843 | **0.001** |
| s (elevational difference) | 6.024 | 48.426 | **0.001** |
| s (hours distance) | 1.353 | 10.637 | **0.043** |
| s (months distance) | 5.800 | 7.876 | **0.045** |
| s (years distance) | 7.135 | 13.007 | **0.020** |
| s (sampling intensity distance) | 1.010 | 5.437 | 0.267 |
| s (methods distance) | 7.878 | 17.094 | **0.003** |

**Table S7. Multiple drivers of plant-frugivore network structural dissimilarity.** Here, we used the binary version of ecoregion and biome distance matrices. *P* values were calculated using a combination of generalized additive models and multiple regression on distance matrices. EDF represents the estimated degrees of freedom for each smooth term in the model. Bold values indicate statistically significant results ($P < 0.05$). *N* pairs of networks = 19,110.

| **Parametric coefficients** | **Estimate** | **t** | **p** |
|---|---|---|---|
| Intercept | 2.689 | 222.572 | **0.002** |
| Ecoregion (same) | 0.043 | 0.632 | 0.788 |
| Biome (same) | -0.028 | -1.345 | 0.770 |
| **Smooth Terms** | **EDF** | **F** | **p** |
| s (local footprint distance) | 5.923 | 9.346 | 0.429 |
| s (spatial distance) | 8.474 | 20.408 | **0.021** |
| s (elevational difference) | 8.220 | 5.509 | 0.749 |
| s (hours distance) | 8.006 | 7.944 | 0.969 |
| s (months distance) | 5.961 | 7.077 | 0.693 |
| s (years distance) | 6.868 | 14.999 | 0.461 |
| s (sampling intensity distance) | 8.762 | 238.987 | **0.002** |
| s (methods distance) | 8.586 | 17.372 | 0.231 |

**Table S8. Multiple drivers of plant-frugivore network structural dissimilarity.** Here, we used the quantitative version (environmental dissimilarity) of ecoregion and biome distance matrices. *P* values were calculated using a combination of generalized additive models and multiple regression on distance matrices. EDF represents the estimated degrees of freedom for each smooth term in the model. Bold values indicate statistically significant results ($P < 0.05$). *N* pairs of networks = 19,110.

| Smooth Terms | EDF | F | p |
|---|---|---|---|
| s (ecoregion distance) | 4.272 | 15.275 | 0.193 |
| s (biome distance) | 7.697 | 12.115 | 0.568 |
| s (local footprint distance) | 5.993 | 9.264 | 0.438 |
| s (spatial distance) | 8.465 | 18.465 | **0.018** |
| s (elevational difference) | 8.290 | 5.679 | 0.713 |
| s (hours distance) | 7.857 | 8.913 | 0.955 |
| s (months distance) | 6.173 | 8.239 | 0.606 |
| s (years distance) | 6.751 | 12.872 | 0.545 |
| s (sampling intensity distance) | 8.760 | 239.475 | **0.002** |
| s (methods distance) | 8.501 | 15.584 | 0.257 |

**Table S9**. **Multiple drivers of species turnover (βs) on plant-frugivore networks.** Here, we used a buffer zone of 500 km and the alternative scenario 1 (see 'Alternative scenarios' section) during the data cleaning process. The binary versions of ecoregion and biome distance matrices were used for estimating the effects of ecoregion and biome borders on the response variable. *P* values were calculated using a combination of generalized additive models and multiple regression on distance matrices. EDF represents the estimated degrees of freedom for each smooth term in the model. Bold values indicate statistically significant results ($P < 0.05$). *N* pairs of networks = 19,110.

| **Parametric coefficients** | **Estimate** | **t** | **p** |
|---|---|---|---|
| Intercept | 0.976 | 1735.325 | **0.001** |
| Ecoregion (same) | -0.122 | -38.147 | **0.001** |
| Biome (same) | -0.008 | -8.809 | **0.001** |
| **Smooth Terms** | **EDF** | **F** | **p** |
| s (local footprint distance) | 8.312 | 28.538 | **0.001** |
| s (spatial distance) | 8.867 | 725.453 | **0.001** |
| s (elevational difference) | 5.600 | 99.711 | **0.001** |
| s (hours distance) | 6.928 | 4.042 | 0.580 |
| s (months distance) | 6.761 | 6.566 | 0.083 |
| s (years distance) | 6.412 | 7.472 | 0.059 |
| s (sampling intensity distance) | 1.001 | 26.885 | **0.005** |
| s (methods distance) | 8.032 | 10.833 | **0.023** |

**Table S10**. **Multiple drivers of species turnover ($\beta_S$) on plant-frugivore networks.** Here, we used a buffer zone of 500 km and the alternative scenario 2 (see 'Alternative scenarios' section) during the data cleaning process. The binary versions of ecoregion and biome distance matrices were used for estimating the effects of ecoregion and biome borders on the response variable. *P* values were calculated using a combination of generalized additive models and multiple regression on distance matrices. EDF represents the estimated degrees of freedom for each smooth term in the model. Bold values indicate statistically significant results ($P < 0.05$). *N* pairs of networks = 19,110.

| **Parametric coefficients** | **Estimate** | **t** | **p** |
|---|---|---|---|
| Intercept | 0.976 | 1752.859 | **0.001** |
| Ecoregion (same) | -0.123 | -38.615 | **0.001** |
| Biome (same) | -0.008 | -8.084 | **0.001** |
| **Smooth Terms** | **EDF** | **F** | **p** |
| s (local footprint distance) | 8.437 | 28.851 | **0.001** |
| s (spatial distance) | 8.865 | 719.288 | **0.001** |
| s (elevational difference) | 5.600 | 99.486 | **0.001** |
| s (hours distance) | 7.126 | 4.330 | 0.559 |
| s (months distance) | 4.001 | 6.532 | 0.091 |
| s (years distance) | 6.548 | 8.206 | 0.069 |
| s (sampling intensity distance) | 3.464 | 8.113 | 0.166 |
| s (methods distance) | 8.114 | 11.641 | **0.013** |

**Table S11**. **Multiple drivers of species turnover ($\beta_S$) on plant-frugivore networks.** Here, we used a buffer zone of 100 km and the alternative scenario 1 (see 'Alternative scenarios' section) during the data cleaning process. The binary versions of ecoregion and biome distance matrices were used for estimating the effects of ecoregion and biome borders on the response variable. *P* values were calculated using a combination of generalized additive models and multiple regression on distance matrices. EDF represents the estimated degrees of freedom for each smooth term in the model. Bold values indicate statistically significant results ($P < 0.05$). *N* pairs of networks = 19,110.

| **Parametric coefficients** | **Estimate** | **t** | **p** |
|---|---|---|---|
| Intercept | 0.976 | 1736.530 | **0.001** |
| Ecoregion (same) | -0.122 | -38.181 | **0.001** |
| Biome (same) | -0.009 | -8.781 | **0.001** |
| **Smooth Terms** | **EDF** | **F** | **p** |
| s (local footprint distance) | 8.317 | 28.664 | **0.002** |
| s (spatial distance) | 8.866 | 725.286 | **0.001** |
| s (elevational difference) | 5.606 | 99.783 | **0.001** |
| s (hours distance) | 6.888 | 3.931 | 0.606 |
| s (months distance) | 6.827 | 6.601 | 0.091 |
| s (years distance) | 6.406 | 7.500 | 0.073 |
| s (sampling intensity distance) | 1.002 | 26.760 | **0.008** |
| s (methods distance) | 8.029 | 10.895 | **0.016** |

**Table S12**. **Multiple drivers of species turnover ($\beta_S$) on plant-frugivore networks.** Here, we used a buffer zone of 100 km and the alternative scenario 2 (see 'Alternative scenarios' section) during the data cleaning process. The binary versions of ecoregion and biome distance matrices were used for estimating the effects of ecoregion and biome borders on the response variable. *P* values were calculated using a combination of generalized additive models and multiple regression on distance matrices. EDF represents the estimated degrees of freedom for each smooth term in the model. Bold values indicate statistically significant results ($P < 0.05$). *N* pairs of networks = 19,110.

| **Parametric coefficients** | **Estimate** | **t** | **p** |
|---|---|---|---|
| Intercept | 0.976 | 1755.726 | **0.001** |
| Ecoregion (same) | -0.122 | -38.561 | **0.001** |
| Biome (same) | -0.008 | -8.354 | **0.001** |
| **Smooth Terms** | **EDF** | **F** | **p** |
| s (local footprint distance) | 8.341 | 29.073 | **0.002** |
| s (spatial distance) | 8.863 | 716.735 | **0.001** |
| s (elevational difference) | 5.578 | 100.041 | **0.001** |
| s (hours distance) | 6.987 | 3.990 | 0.592 |
| s (months distance) | 6.819 | 6.693 | 0.107 |
| s (years distance) | 6.484 | 7.966 | 0.063 |
| s (sampling intensity distance) | 1.000 | 24.580 | **0.005** |
| s (methods distance) | 8.013 | 11.066 | **0.018** |

**Table S13. Multiple drivers of species turnover ($\beta_S$) on plant-frugivore networks.** Here, we used a buffer zone of 100 km and the alternative scenario 3 (see 'Alternative scenarios' section) during the data cleaning process. The binary versions of ecoregion and biome distance matrices were used for estimating the effects of ecoregion and biome borders on the response variable. *P* values were calculated using a combination of generalized additive models and multiple regression on distance matrices. EDF represents the estimated degrees of freedom for each smooth term in the model. Bold values indicate statistically significant results ($P < 0.05$). *N* pairs of networks = 19,110.

| **Parametric coefficients** | **Estimate** | **t** | **p** |
|---|---|---|---|
| Intercept | 0.976 | 1735.345 | **0.001** |
| Ecoregion (same) | -0.122 | -38.157 | **0.001** |
| Biome (same) | -0.009 | -8.775 | **0.001** |
| **Smooth Terms** | **EDF** | **F** | **p** |
| s (local footprint distance) | 8.317 | 28.665 | **0.002** |
| s (spatial distance) | 8.866 | 723.914 | **0.001** |
| s (elevational difference) | 5.587 | 99.935 | **0.001** |
| s (hours distance) | 6.918 | 4.014 | 0.605 |
| s (months distance) | 6.783 | 6.589 | 0.100 |
| s (years distance) | 6.406 | 7.492 | 0.078 |
| s (sampling intensity distance) | 1.000 | 26.866 | **0.004** |
| s (methods distance) | 8.033 | 10.910 | **0.011** |

**Table S14**. **Multiple drivers of species turnover ($\beta_S$) on plant-frugivore networks.** Here, we used a buffer zone of 1000 km and the alternative scenario 1 (see 'Alternative scenarios' section) during the data cleaning process. The binary versions of ecoregion and biome distance matrices were used for estimating the effects of ecoregion and biome borders on the response variable. *P* values were calculated using a combination of generalized additive models and multiple regression on distance matrices. EDF represents the estimated degrees of freedom for each smooth term in the model. Bold values indicate statistically significant results ($P < 0.05$). *N* pairs of networks = 19,110.

| **Parametric coefficients** | **Estimate** | **t** | **p** |
|---|---|---|---|
| Intercept | 0.976 | 1734.871 | **0.001** |
| Ecoregion (same) | -0.122 | -38.147 | **0.001** |
| Biome (same) | -0.009 | -8.789 | **0.001** |
| **Smooth Terms** | **EDF** | **F** | **p** |
| s (local footprint distance) | 8.309 | 28.531 | **0.001** |
| s (spatial distance) | 8.866 | 725.141 | **0.001** |
| s (elevational difference) | 5.605 | 99.321 | **0.001** |
| s (hours distance) | 6.911 | 4.049 | 0.602 |
| s (months distance) | 6.761 | 6.579 | 0.099 |
| s (years distance) | 6.414 | 7.440 | 0.067 |
| s (sampling intensity distance) | 1.002 | 26.590 | **0.005** |
| s (methods distance) | 8.030 | 10.869 | **0.019** |

**Table S15**. **Multiple drivers of species turnover ($\beta_S$) on plant-frugivore networks.** Here, we used a buffer zone of 1000 km and the alternative scenario 2 (see 'Alternative scenarios' section) during the data cleaning process. The binary versions of ecoregion and biome distance matrices were used for estimating the effects of ecoregion and biome borders on the response variable. *P* values were calculated using a combination of generalized additive models and multiple regression on distance matrices. EDF represents the estimated degrees of freedom for each smooth term in the model. Bold values indicate statistically significant results ($P < 0.05$). *N* pairs of networks = 19,110.

| **Parametric coefficients** | **Estimate** | **t** | **p** |
|---|---|---|---|
| Intercept | 0.976 | 1755.726 | **0.001** |
| Ecoregion (same) | -0.122 | -38.561 | **0.001** |
| Biome (same) | -0.008 | -8.354 | **0.001** |
| **Smooth Terms** | **EDF** | **F** | **p** |
| s (local footprint distance) | 8.341 | 29.073 | **0.001** |
| s (spatial distance) | 8.863 | 716.735 | **0.001** |
| s (elevational difference) | 5.578 | 100.041 | **0.001** |
| s (hours distance) | 6.987 | 3.990 | 0.608 |
| s (months distance) | 6.819 | 6.693 | 0.087 |
| s (years distance) | 6.484 | 7.966 | 0.061 |
| s (sampling intensity distance) | 1.000 | 24.580 | **0.008** |
| s (methods distance) | 8.013 | 11.066 | **0.016** |

**Table S16**. **Multiple drivers of species turnover ($\beta_S$) on plant-frugivore networks.** Here, we used a buffer zone of 1000 km and the alternative scenario 3 (see 'Alternative scenarios' section) during the data cleaning process. The binary versions of ecoregion and biome distance matrices were used for estimating the effects of ecoregion and biome borders on the response variable. *P* values were calculated using a combination of generalized additive models and multiple regression on distance matrices. EDF represents the estimated degrees of freedom for each smooth term in the model. Bold values indicate statistically significant results ($P < 0.05$). *N* pairs of networks = 19,110.

| **Parametric coefficients** | **Estimate** | **t** | **p** |
|---|---|---|---|
| Intercept | 0.976 | 1733.860 | **0.001** |
| Ecoregion (same) | -0.122 | -38.095 | **0.001** |
| Biome (same) | -0.009 | -8.778 | **0.001** |
| **Smooth Terms** | **EDF** | **F** | **p** |
| s (local footprint distance) | 8.308 | 28.497 | **0.001** |
| s (spatial distance) | 8.866 | 725.333 | **0.001** |
| s (elevational difference) | 5.594 | 99.561 | **0.001** |
| s (hours distance) | 6.899 | 4.011 | 0.608 |
| s (months distance) | 6.744 | 6.537 | 0.109 |
| s (years distance) | 6.404 | 7.389 | 0.063 |
| s (sampling intensity distance) | 1.004 | 26.506 | **0.006** |
| s (methods distance) | 8.037 | 10.951 | **0.021** |

**Table S17**. **Multiple drivers of plant-frugivore interaction dissimilarity ($\beta_{WN}$).** Here, we used a buffer zone of 500 km and the alternative scenario 1 (see 'Alternative scenarios' section) during the data cleaning process. The binary versions of ecoregion and biome distance matrices were used for estimating the effects of ecoregion and biome borders on the response variable. *P* values were calculated using a combination of generalized additive models and multiple regression on distance matrices. EDF represents the estimated degrees of freedom for each smooth term in the model. Bold values indicate statistically significant results ($P < 0.05$). *N* pairs of networks = 19,110.

| **Parametric coefficients** | **Estimate** | **t** | **p** |
|---|---|---|---|
| Intercept | 0.997 | 2966.347 | **0.001** |
| Ecoregion (same) | -0.070 | -36.417 | **0.001** |
| Biome (same) | -0.002 | -3.317 | **0.039** |
| **Smooth Terms** | **EDF** | **F** | **p** |
| s (local footprint distance) | 8.536 | 30.035 | **0.001** |
| s (spatial distance) | 8.785 | 65.220 | **0.001** |
| s (elevational difference) | 6.185 | 47.606 | **0.001** |
| s (hours distance) | 1.545 | 5.545 | 0.294 |
| s (months distance) | 5.502 | 6.966 | 0.074 |
| s (years distance) | 7.216 | 11.880 | **0.013** |
| s (sampling intensity distance) | 1.062 | 4.686 | 0.331 |
| s (methods distance) | 7.848 | 15.987 | **0.004** |

**Table S18**. **Multiple drivers of plant-frugivore interaction dissimilarity ($\beta_{WN}$).** Here, we used a buffer zone of 500 km and the alternative scenario 2 (see 'Alternative scenarios' section) during the data cleaning process. The binary versions of ecoregion and biome distance matrices were used for estimating the effects of ecoregion and biome borders on the response variable. *P* values were calculated using a combination of generalized additive models and multiple regression on distance matrices. EDF represents the estimated degrees of freedom for each smooth term in the model. Bold values indicate statistically significant results ($P < 0.05$). *N* pairs of networks = 19,110.

| **Parametric coefficients** | **Estimate** | **t** | **p** |
|---|---|---|---|
| Intercept | 0.997 | 3002.392 | **0.001** |
| Ecoregion (same) | -0.069 | -36.473 | **0.001** |
| Biome (same) | -0.002 | -3.313 | **0.034** |
| **Smooth Terms** | **EDF** | **F** | **p** |
| s (local footprint distance) | 8.551 | 30.504 | **0.001** |
| s (spatial distance) | 8.783 | 64.233 | **0.001** |
| s (elevational difference) | 6.107 | 47.553 | **0.001** |
| s (hours distance) | 1.590 | 5.325 | 0.307 |
| s (months distance) | 5.475 | 7.030 | 0.092 |
| s (years distance) | 7.216 | 11.941 | **0.022** |
| s (sampling intensity distance) | 1.003 | 5.041 | 0.319 |
| s (methods distance) | 7.867 | 16.082 | **0.003** |

**Table S19**. **Multiple drivers of plant-frugivore interaction dissimilarity ($\beta_{WN}$).** Here, we used a buffer zone of 100 km and the alternative scenario 1 (see 'Alternative scenarios' section) during the data cleaning process. The binary versions of ecoregion and biome distance matrices were used for estimating the effects of ecoregion and biome borders on the response variable. *P* values were calculated using a combination of generalized additive models and multiple regression on distance matrices. EDF represents the estimated degrees of freedom for each smooth term in the model. Bold values indicate statistically significant results ($P < 0.05$). *N* pairs of networks = 19,110.

| **Parametric coefficients** | **Estimate** | **t** | **p** |
|---|---|---|---|
| Intercept | 0.997 | 2966.503 | **0.001** |
| Ecoregion (same) | -0.070 | -36.418 | **0.001** |
| Biome (same) | -0.002 | -3.321 | **0.047** |
| **Smooth Terms** | **EDF** | **F** | **p** |
| s (local footprint distance) | 8.536 | 30.011 | **0.001** |
| s (spatial distance) | 8.785 | 65.161 | **0.001** |
| s (elevational difference) | 6.190 | 47.625 | **0.001** |
| s (hours distance) | 1.546 | 5.546 | 0.272 |
| s (months distance) | 5.504 | 6.965 | 0.074 |
| s (years distance) | 7.215 | 11.883 | **0.021** |
| s (sampling intensity distance) | 1.056 | 4.744 | 0.330 |
| s (methods distance) | 7.851 | 16.023 | **0.005** |

**Table S20**. **Multiple drivers of plant-frugivore interaction dissimilarity ($\beta_{WN}$).** Here, we used a buffer zone of 100 km and the alternative scenario 2 (see ‘Alternative scenarios’ section) during the data cleaning process. The binary versions of ecoregion and biome distance matrices were used for estimating the effects of ecoregion and biome borders on the response variable. *P* values were calculated using a combination of generalized additive models and multiple regression on distance matrices. EDF represents the estimated degrees of freedom for each smooth term in the model. Bold values indicate statistically significant results ($P < 0.05$). *N* pairs of networks = 19,110.

| **Parametric coefficients** | **Estimate** | **t** | **p** |
|---|---|---|---|
| Intercept | 0.997 | 3002.382 | **0.001** |
| Ecoregion (same) | -0.069 | -36.474 | **0.001** |
| Biome (same) | -0.002 | -3.312 | **0.049** |
| **Smooth Terms** | **EDF** | **F** | **p** |
| s (local footprint distance) | 8.551 | 30.506 | **0.002** |
| s (spatial distance) | 8.782 | 64.153 | **0.001** |
| s (elevational difference) | 6.109 | 47.538 | **0.001** |
| s (hours distance) | 1.579 | 5.376 | 0.298 |
| s (months distance) | 5.483 | 7.037 | 0.075 |
| s (years distance) | 7.217 | 11.954 | **0.019** |
| s (sampling intensity distance) | 1.003 | 5.036 | 0.311 |
| s (methods distance) | 7.867 | 16.089 | **0.004** |

**Table S21. Multiple drivers of plant-frugivore interaction dissimilarity ($\beta_{WN}$).** Here, we used a buffer zone of 100 km and the alternative scenario 3 (see 'Alternative scenarios' section) during the data cleaning process. The binary versions of ecoregion and biome distance matrices were used for estimating the effects of ecoregion and biome borders on the response variable. *P* values were calculated using a combination of generalized additive models and multiple regression on distance matrices. EDF represents the estimated degrees of freedom for each smooth term in the model. Bold values indicate statistically significant results ($P < 0.05$). *N* pairs of networks = 19,110.

| **Parametric coefficients** | **Estimate** | **t** | **p** |
|---|---|---|---|
| Intercept | 0.997 | 2964.236 | **0.001** |
| Ecoregion (same) | -0.070 | -36.405 | **0.001** |
| Biome (same) | -0.002 | -3.324 | **0.046** |
| **Smooth Terms** | **EDF** | **F** | **p** |
| s (local footprint distance) | 8.534 | 29.980 | **0.001** |
| s (spatial distance) | 8.785 | 65.228 | **0.001** |
| s (elevational difference) | 6.171 | 47.691 | **0.001** |
| s (hours distance) | 1.559 | 5.453 | 0.301 |
| s (months distance) | 5.490 | 6.908 | 0.076 |
| s (years distance) | 7.210 | 11.881 | **0.020** |
| s (sampling intensity distance) | 1.022 | 5.148 | 0.281 |
| s (methods distance) | 7.850 | 16.024 | **0.004** |

**Table S22. Multiple drivers of plant-frugivore interaction dissimilarity ($\beta_{WN}$).** Here, we used a buffer zone of 1000 km and the alternative scenario 1 (see 'Alternative scenarios' section) during the data cleaning process. The binary versions of ecoregion and biome distance matrices were used for estimating the effects of ecoregion and biome borders on the response variable. *P* values were calculated using a combination of generalized additive models and multiple regression on distance matrices. EDF represents the estimated degrees of freedom for each smooth term in the model. Bold values indicate statistically significant results ($P < 0.05$). *N* pairs of networks = 19,110.

| **Parametric coefficients** | **Estimate** | **t** | **p** |
|---|---|---|---|
| Intercept | 0.997 | 2966.167 | **0.001** |
| Ecoregion (same) | -0.070 | -36.419 | **0.001** |
| Biome (same) | -0.002 | -3.311 | **0.032** |
| **Smooth Terms** | **EDF** | **F** | **p** |
| s (local footprint distance) | 8.536 | 30.036 | **0.001** |
| s (spatial distance) | 8.785 | 65.100 | **0.001** |
| s (elevational difference) | 6.187 | 47.586 | **0.001** |
| s (hours distance) | 1.532 | 5.585 | 0.299 |
| s (months distance) | 5.511 | 6.974 | 0.076 |
| s (years distance) | 7.217 | 11.890 | **0.019** |
| s (sampling intensity distance) | 1.085 | 4.382 | 0.377 |
| s (methods distance) | 7.849 | 15.996 | **0.004** |

**Table S23. Multiple drivers of plant-frugivore interaction dissimilarity ($\beta_{WN}$).** Here, we used a buffer zone of 1000 km and the alternative scenario 2 (see 'Alternative scenarios' section) during the data cleaning process. The binary versions of ecoregion and biome distance matrices were used for estimating the effects of ecoregion and biome borders on the response variable. *P* values were calculated using a combination of generalized additive models and multiple regression on distance matrices. EDF represents the estimated degrees of freedom for each smooth term in the model. Bold values indicate statistically significant results ($P < 0.05$). *N* pairs of networks = 19,110.

| **Parametric coefficients** | **Estimate** | **t** | **p** |
|---|---|---|---|
| Intercept | 0.997 | 3002.382 | **0.001** |
| Ecoregion (same) | -0.069 | -36.474 | **0.001** |
| Biome (same) | -0.002 | -3.312 | **0.048** |
| **Smooth Terms** | **EDF** | **F** | **p** |
| s (local footprint distance) | 8.551 | 30.506 | **0.002** |
| s (spatial distance) | 8.782 | 64.153 | **0.001** |
| s (elevational difference) | 6.109 | 47.538 | **0.001** |
| s (hours distance) | 1.579 | 5.376 | 0.311 |
| s (months distance) | 5.483 | 7.037 | 0.054 |
| s (years distance) | 7.217 | 11.954 | **0.017** |
| s (sampling intensity distance) | 1.003 | 5.036 | 0.320 |
| s (methods distance) | 7.867 | 16.089 | **0.004** |

**Table S24**. **Multiple drivers of plant-frugivore interaction dissimilarity ($\beta_{WN}$).** Here, we used a buffer zone of 1000 km and the alternative scenario 3 (see 'Alternative scenarios' section) during the data cleaning process. The binary versions of ecoregion and biome distance matrices were used for estimating the effects of ecoregion and biome borders on the response variable. *P* values were calculated using a combination of generalized additive models and multiple regression on distance matrices. EDF represents the estimated degrees of freedom for each smooth term in the model. Bold values indicate statistically significant results ($P < 0.05$). *N* pairs of networks = 19,110.

| **Parametric coefficients** | **Estimate** | **t** | **p** |
|---|---|---|---|
| Intercept | 0.997 | 2964.095 | **0.001** |
| Ecoregion (same) | -0.070 | -36.404 | **0.001** |
| Biome (same) | -0.002 | -3.318 | **0.042** |
| **Smooth Terms** | **EDF** | **F** | **p** |
| s (local footprint distance) | 8.534 | 29.989 | **0.002** |
| s (spatial distance) | 8.785 | 65.276 | **0.001** |
| s (elevational difference) | 6.170 | 47.687 | **0.001** |
| s (hours distance) | 1.547 | 5.482 | 0.300 |
| s (months distance) | 5.491 | 6.909 | 0.073 |
| s (years distance) | 7.210 | 11.857 | **0.020** |
| s (sampling intensity distance) | 1.026 | 4.983 | 0.287 |
| s (methods distance) | 7.849 | 16.010 | **0.003** |

**Table S25**. **Multiple drivers of plant-frugivore network structural dissimilarity.** Here, we used a buffer zone of 500 km and the alternative scenario 1 (see 'Alternative scenarios' section) during the data cleaning process. The binary versions of ecoregion and biome distance matrices were used for estimating the effects of ecoregion and biome borders on the response variable. *P* values were calculated using a combination of generalized additive models and multiple regression on distance matrices. EDF represents the estimated degrees of freedom for each smooth term in the model. Bold values indicate statistically significant results ($P < 0.05$). *N* pairs of networks = 19,110.

| **Parametric coefficients** | **Estimate** | **t** | **p** |
|---|---|---|---|
| Intercept | 2.686 | 221.962 | **0.004** |
| Ecoregion (same) | 0.044 | 0.646 | 0.775 |
| Biome (same) | -0.024 | -1.115 | 0.826 |
| **Smooth Terms** | **EDF** | **F** | **p** |
| s (local footprint distance) | 5.948 | 9.481 | 0.439 |
| s (spatial distance) | 8.473 | 20.322 | **0.015** |
| s (elevational difference) | 8.233 | 5.501 | 0.724 |
| s (hours distance) | 8.051 | 7.960 | 0.968 |
| s (months distance) | 6.239 | 7.217 | 0.667 |
| s (years distance) | 6.830 | 13.941 | 0.497 |
| s (sampling intensity distance) | 8.759 | 240.837 | **0.001** |
| s (methods distance) | 8.595 | 17.496 | 0.233 |

**Table S26. Multiple drivers of plant-frugivore network structural dissimilarity.** Here, we used a buffer zone of 500 km and the alternative scenario 2 (see 'Alternative scenarios' section) during the data cleaning process. The binary versions of ecoregion and biome distance matrices were used for estimating the effects of ecoregion and biome borders on the response variable. *P* values were calculated using a combination of generalized additive models and multiple regression on distance matrices. EDF represents the estimated degrees of freedom for each smooth term in the model. Bold values indicate statistically significant results ($P < 0.05$). *N* pairs of networks = 19,110.

| **Parametric coefficients** | **Estimate** | **t** | **p** |
|---|---|---|---|
| Intercept | 2.685 | 222.539 | **0.002** |
| Ecoregion (same) | 0.084 | 1.229 | 0.561 |
| Biome (same) | -0.024 | -1.157 | 0.801 |
| **Smooth Terms** | **EDF** | **F** | **p** |
| s (local footprint distance) | 5.417 | 9.472 | 0.460 |
| s (spatial distance) | 8.587 | 28.061 | **0.002** |
| s (elevational difference) | 7.800 | 3.418 | 0.904 |
| s (hours distance) | 8.088 | 7.568 | 0.973 |
| s (months distance) | 7.129 | 7.330 | 0.682 |
| s (years distance) | 6.823 | 12.437 | 0.555 |
| s (sampling intensity distance) | 8.758 | 275.291 | **0.001** |
| s (methods distance) | 8.550 | 18.139 | 0.191 |

**Table S27. Multiple drivers of plant-frugivore network structural dissimilarity.** Here, we used a buffer zone of 100 km and the alternative scenario 1 (see 'Alternative scenarios' section) during the data cleaning process. The binary versions of ecoregion and biome distance matrices were used for estimating the effects of ecoregion and biome borders on the response variable. *P* values were calculated using a combination of generalized additive models and multiple regression on distance matrices. EDF represents the estimated degrees of freedom for each smooth term in the model. Bold values indicate statistically significant results ($P < 0.05$). *N* pairs of networks = 19,110.

| **Parametric coefficients** | **Estimate** | **t** | **p** |
|---|---|---|---|
| Intercept | 2.691 | 222.709 | **0.007** |
| Ecoregion (same) | 0.052 | 0.757 | 0.743 |
| Biome (same) | -0.028 | -1.364 | 0.762 |
| **Smooth Terms** | **EDF** | **F** | **p** |
| s (local footprint distance) | 5.834 | 9.562 | 0.428 |
| s (spatial distance) | 8.470 | 20.654 | **0.018** |
| s (elevational difference) | 8.080 | 4.412 | 0.817 |
| s (hours distance) | 8.130 | 8.456 | 0.965 |
| s (months distance) | 6.321 | 7.283 | 0.647 |
| s (years distance) | 6.827 | 13.789 | 0.501 |
| s (sampling intensity distance) | 8.745 | 241.194 | **0.003** |
| s (methods distance) | 8.590 | 17.524 | 0.209 |

**Table S28**. **Multiple drivers of plant-frugivore network structural dissimilarity.** Here, we used a buffer zone of 100 km and the alternative scenario 2 (see 'Alternative scenarios' section) during the data cleaning process. The binary versions of ecoregion and biome distance matrices were used for estimating the effects of ecoregion and biome borders on the response variable. *P* values were calculated using a combination of generalized additive models and multiple regression on distance matrices. EDF represents the estimated degrees of freedom for each smooth term in the model. Bold values indicate statistically significant results ($P < 0.05$). *N* pairs of networks = 19,110.

| **Parametric coefficients** | **Estimate** | **t** | **p** |
|---|---|---|---|
| Intercept | 2.684 | 222.432 | **0.004** |
| Ecoregion (same) | 0.089 | 1.311 | 0.549 |
| Biome (same) | -0.023 | -1.085 | 0.812 |
| **Smooth Terms** | **EDF** | **F** | **p** |
| s (local footprint distance) | 5.330 | 9.475 | 0.436 |
| s (spatial distance) | 8.590 | 28.764 | **0.003** |
| s (elevational difference) | 1.026 | 4.544 | 0.803 |
| s (hours distance) | 8.122 | 7.758 | 0.981 |
| s (months distance) | 7.189 | 7.442 | 0.677 |
| s (years distance) | 6.821 | 12.365 | 0.583 |
| s (sampling intensity distance) | 8.761 | 275.772 | **0.001** |
| s (methods distance) | 8.540 | 17.893 | 0.205 |

**Table S29**. **Multiple drivers of plant-frugivore network structural dissimilarity.** Here, we used a buffer zone of 100 km and the alternative scenario 3 (see 'Alternative scenarios' section) during the data cleaning process. The binary versions of ecoregion and biome distance matrices were used for estimating the effects of ecoregion and biome borders on the response variable. *P* values were calculated using a combination of generalized additive models and multiple regression on distance matrices. EDF represents the estimated degrees of freedom for each smooth term in the model. Bold values indicate statistically significant results ($P < 0.05$). *N* pairs of networks = 19,110.

| **Parametric coefficients** | **Estimate** | **t** | **p** |
|---|---|---|---|
| Intercept | 2.689 | 222.557 | **0.008** |
| Ecoregion (same) | 0.044 | 0.639 | 0.754 |
| Biome (same) | -0.031 | -1.443 | 0.741 |
| **Smooth Terms** | **EDF** | **F** | **p** |
| s (local footprint distance) | 5.869 | 9.131 | 0.446 |
| s (spatial distance) | 8.479 | 20.589 | **0.021** |
| s (elevational difference) | 8.217 | 5.476 | 0.755 |
| s (hours distance) | 8.052 | 7.939 | 0.966 |
| s (months distance) | 6.005 | 7.020 | 0.675 |
| s (years distance) | 6.834 | 14.956 | 0.411 |
| s (sampling intensity distance) | 8.746 | 238.220 | **0.003** |
| s (methods distance) | 8.583 | 17.496 | 0.206 |

**Table S30**. **Multiple drivers of plant-frugivore network structural dissimilarity.** Here, we used a buffer zone of 1000 km and the alternative scenario 1 (see 'Alternative scenarios' section) during the data cleaning process. The binary versions of ecoregion and biome distance matrices were used for estimating the effects of ecoregion and biome borders on the response variable. *P* values were calculated using a combination of generalized additive models and multiple regression on distance matrices. EDF represents the estimated degrees of freedom for each smooth term in the model. Bold values indicate statistically significant results ($P < 0.05$). *N* pairs of networks = 19,110.

| **Parametric coefficients** | **Estimate** | **t** | **p** |
|---|---|---|---|
| Intercept | 2.687 | 222.335 | **0.001** |
| Ecoregion (same) | 0.047 | 0.681 | 0.776 |
| Biome (same) | -0.026 | -1.251 | 0.802 |
| **Smooth Terms** | **EDF** | **F** | **p** |
| s (local footprint distance) | 5.954 | 9.761 | 0.432 |
| s (spatial distance) | 8.483 | 20.514 | **0.010** |
| s (elevational difference) | 8.243 | 5.492 | 0.736 |
| s (hours distance) | 8.009 | 7.896 | 0.970 |
| s (months distance) | 6.128 | 6.943 | 0.699 |
| s (years distance) | 6.852 | 13.832 | 0.496 |
| s (sampling intensity distance) | 8.789 | 245.694 | **0.002** |
| s (methods distance) | 8.593 | 17.437 | 0.229 |

**Table S31**. **Multiple drivers of plant-frugivore network structural dissimilarity.** Here, we used a buffer zone of 1000 km and the alternative scenario 2 (see 'Alternative scenarios' section) during the data cleaning process. The binary versions of ecoregion and biome distance matrices were used for estimating the effects of ecoregion and biome borders on the response variable. *P* values were calculated using a combination of generalized additive models and multiple regression on distance matrices. EDF represents the estimated degrees of freedom for each smooth term in the model. Bold values indicate statistically significant results ($P < 0.05$). *N* pairs of networks = 19,110.

| **Parametric coefficients** | **Estimate** | **t** | **p** |
|---|---|---|---|
| Intercept | 2.685 | 222.527 | **0.004** |
| Ecoregion (same) | 0.084 | 1.225 | 0.562 |
| Biome (same) | -0.022 | -1.058 | 0.844 |
| **Smooth Terms** | **EDF** | **F** | **p** |
| s (local footprint distance) | 5.417 | 9.454 | 0.427 |
| s (spatial distance) | 8.588 | 28.139 | **0.008** |
| s (elevational difference) | 7.796 | 3.409 | 0.893 |
| s (hours distance) | 8.098 | 7.547 | 0.977 |
| s (months distance) | 7.123 | 7.341 | 0.669 |
| s (years distance) | 6.851 | 12.533 | 0.570 |
| s (sampling intensity distance) | 8.757 | 275.296 | **0.001** |
| s (methods distance) | 8.551 | 18.041 | 0.182 |

**Table S33. Multiple drivers of plant-frugivore network structural dissimilarity.** Here, we used a buffer zone of 1000 km and the alternative scenario 3 (see 'Alternative scenarios' section) during the data cleaning process. The binary versions of ecoregion and biome distance matrices were used for estimating the effects of ecoregion and biome borders on the response variable. *P* values were calculated using a combination of generalized additive models and multiple regression on distance matrices. EDF represents the estimated degrees of freedom for each smooth term in the model. Bold values indicate statistically significant results ($P < 0.05$). *N* pairs of networks = 19,110.

| **Parametric coefficients** | **Estimate** | **t** | **p** |
|---|---|---|---|
| Intercept | 2.692 | 223.088 | **0.008** |
| Ecoregion (same) | 0.045 | 0.663 | 0.766 |
| Biome (same) | -0.033 | -1.581 | 0.748 |
| **Smooth Terms** | **EDF** | **F** | **p** |
| s (local footprint distance) | 5.943 | 9.649 | 0.423 |
| s (spatial distance) | 8.491 | 20.649 | **0.013** |
| s (elevational difference) | 8.230 | 5.556 | 0.727 |
| s (hours distance) | 8.063 | 8.161 | 0.956 |
| s (months distance) | 5.980 | 6.955 | 0.711 |
| s (years distance) | 6.778 | 14.670 | 0.479 |
| s (sampling intensity distance) | 8.792 | 243.787 | **0.001** |
| s (methods distance) | 8.578 | 17.155 | 0.237 |

**Table S34. Multiple drivers of plant-frugivore interaction dissimilarity ($\beta_{WN}$).** Here, we used the binary version of ecoregion and biome distance matrices and removed the study with the greatest number of networks in our dataset [study 76 (*79*)] from the data. *P* values were calculated using a combination of generalized additive models and multiple regression on distance matrices. EDF represents the estimated degrees of freedom for each smooth term in the model. Bold values indicate statistically significant results ($P < 0.05$). *N* pairs of networks = 12,880.

| **Parametric coefficients** | **Estimate** | **t** | **p** |
|---|---|---|---|
| Intercept | 0.995 | 2816.925 | **0.001** |
| Ecoregion (same) | -0.077 | -33.132 | **0.001** |
| Biome (same) | -0.0008 | -1.254 | 0.380 |
| **Smooth Terms** | **EDF** | **F** | **p** |
| s (local footprint distance) | 6.871 | 11.919 | **0.005** |
| s (spatial distance) | 8.917 | 139.693 | **0.001** |
| s (elevational difference) | 5.502 | 9.025 | **0.035** |
| s (hours distance) | 2.007 | 7.295 | 0.106 |
| s (months distance) | 7.806 | 23.758 | **0.001** |
| s (years distance) | 8.500 | 33.731 | **0.001** |
| s (sampling intensity distance) | 1.002 | 0.015 | 0.992 |
| s (methods distance) | 8.571 | 61.413 | **0.001** |

**Table S35. Multiple drivers of plant-frugivore network structural dissimilarity.** Here, we used the binary version of ecoregion and biome distance matrices and removed the study with the greatest number of networks in our dataset [study 76 (*79*)] from the data. P values were calculated using a combination of generalized additive models and multiple regression on distance matrices. EDF represents the estimated degrees of freedom for each smooth term in the model. Bold values indicate statistically significant results ($P < 0.05$). $N$ pairs of networks = 12,880.

| **Parametric coefficients** | **Estimate** | **t** | **p** |
|---|---|---|---|
| Intercept | 2.568 | 184.419 | **0.022** |
| Ecoregion (same) | -0.075 | -0.826 | 0.544 |
| Biome (same) | 0.041 | 1.595 | 0.679 |
| **Smooth Terms** | **EDF** | **F** | **p** |
| s (local footprint distance) | 4.419 | 13.240 | 0.121 |
| s (spatial distance) | 8.540 | 27.067 | **0.005** |
| s (elevational difference) | 7.486 | 11.064 | 0.364 |
| s (hours distance) | 7.717 | 8.123 | 0.923 |
| s (months distance) | 6.900 | 5.378 | 0.714 |
| s (years distance) | 5.505 | 10.312 | 0.424 |
| s (sampling intensity distance) | 8.534 | 126.502 | **0.008** |
| s (methods distance) | 8.489 | 14.492 | 0.190 |

**Table S36. Multiple drivers of interaction rewiring ($\beta_{OS}$) on plant-frugivore networks.** Here, we show the results from a generalized additive mixed-effects model (GAMM) using network IDs as random effects (one random factor for each of the pairs across which distance is compared) to account for the non-independence of distances (see 'Rewiring analysis' section). EDF represents the estimated degrees of freedom for each smooth term in the model. Bold values indicate statistically significant results ($P < 0.05$). $N$ pairs of networks = 1,314.

| **Parametric coefficients** | **Estimate** | **t** | **p** |
|---|---|---|---|
| Intercept | 0.576 | 25.267 | **<2e-16** |
| Ecoregion (same) | -0.017 | -0.640 | 0.522 |
| Biome (same) | -0.033 | -1.355 | 0.176 |
| **Smooth Terms** | **EDF** | **F** | **p** |
| s (local footprint distance) | 1.875 | 5.046 | **0.005** |
| s (spatial distance) | 2.869 | 17.933 | **1.62e-10** |
| s (elevational difference) | 3.108 | 13.441 | **7.64e-09** |
| s (hours distance) | 1.001 | 5.624 | 0.018 |
| s (months distance) | 3.431 | 1.439 | 0.140 |
| s (years distance) | 1.000 | 0.907 | 0.341 |
| s (sampling intensity distance) | 2.029 | 3.765 | **0.023** |
| s (methods distance) | 1.000 | 5.126 | **0.024** |

**Table S37. The effect of large-scale ecological boundaries on interaction rewiring ($\beta_{OS}$).** Here, we show the results from a generalized additive mixed-effects model (GAMM) using ecoregion and biome distance metrics as predictors and network IDs as random effects (one random factor for each of the pairs across which distance is compared) to account for the non-independence of distances (see 'Rewiring analysis' section). Bold values indicate statistically significant results ($P < 0.05$). Note that, contrary to the full model (Table S36), the effect of ecoregion boundaries is significant, likely because of their collinearity with our other predictor variables. *N* pairs of networks = 1,314.

| Parametric coefficients | Estimate | t | p |
|---|---|---|---|
| Intercept | 0.579 | 22.975 | **<2e-16** |
| Ecoregion (same) | -0.155 | -6.756 | **2.13e-11** |
| Biome (same) | 0.005 | 0.202 | 0.84 |
